# Parallel enterocyte caspase and cytokine responses couple hypoxia to reoxygenation-dependent intestinal regeneration in *Drosophila*

**DOI:** 10.64898/2026.11.12.766219

**Authors:** Prajakta Bodkhe, Kate Ding, Savraj S Grewal

## Abstract

Intestinal oxygen availability fluctuates during both normal physiology and disease, yet how the intestinal epithelium adapts to these changing environments remains poorly understood. Using the adult *Drosophila* intestine, we show that hypoxia and reoxygenation drive distinct phases of an epithelial damage and repair response. Hypoxia induces two parallel responses in largely separate enterocyte populations: an AMPK-p53-dependent response that primes enterocytes for apoptosis, and an independent Upd cytokine response that activates JAK-STAT signaling in intestinal stem cells (ISCs). Despite activation of both pathways during hypoxia, neither apoptosis nor ISC proliferation occurs until oxygen is restored. Upon reoxygenation, enterocyte loss and ISC proliferation proceed together and both are required for recovery of intestinal length. We further show that HIF-1α/Sima is dispensable in enterocytes but required in progenitor cells for reoxygenation-induced proliferation. Finally, repeated episodes of hypoxia-reoxygenation shorten lifespan, and this is exacerbated by enterocyte p53 knockdown. Together, these findings indicate that the epithelium adapts to fluctuating oxygen by coordinating damage and repair responses in space and time.

## Introduction

The intestinal epithelium is one of the most rapidly renewing tissues in the body. This continuous self-renewal replaces epithelial cells lost to normal wear and to environmental insults. Renewal is sustained by a resident population of intestinal stem cells (ISCs) whose proliferation and differentiation are dynamically coupled to the demand for cell replacement, a process conserved across animal species (Beumer & Clevers, 2021; Biteau *et al*, 2011; Jiang & Edgar, 2012; Li & Jasper, 2016; Micchelli & Perrimon, 2006; Takashima & Hartenstein, 2012; Trubin *et al*, 2024; Wang & Hou, 2010; Zwick *et al*, 2019). How this coupling is achieved and maintained in the face of diverse and changing conditions within the gut environment is a central question in intestinal biology.

One key environmental challenge influencing gut physiology is fluctuating oxygen availability. Although ambient air contains approximately 21% oxygen, oxygen levels within the intestine are substantially lower and vary markedly across intestinal regions. Along the longitudinal axis of the gut, oxygen levels decline from approximately 5% in the duodenum to less than 1% in the colonic lumen (Colgan *et al*, 2016; Solanki & Shah, 2024). A steep oxygen gradient also exists across the intestinal epithelium. In the small intestine, crypt cells located near the vasculature are exposed to higher oxygen levels than cells at the villus tips, which face the near-anoxic gut lumen (Singhal & Shah, 2020; Solanki & Shah, 2024). These low-oxygen states, referred to as physiological hypoxia or physoxia, are not static. For example, oxygen availability can fluctuate during feeding-fasting cycles, as intestinal blood flow, and consequently oxygen delivery, can increase several fold following a meal (Colgan & Taylor, 2010). Oxygen levels are also shaped by the gut microbiota, whose metabolic activity consumes oxygen and further contributes to oxygen heterogeneity within the intestinal environment (Ghoshal *et al*, 2026; Kelly *et al*, 2015; Litvak *et al*, 2018; Zheng *et al*, 2015).

Oxygen delivery to the intestine is also disrupted in a number of different pathologies. For example, in inflammatory bowel diseases such as colitis, local vascular dysfunction and the increased metabolic demand of infiltrating immune cells produce severe hypoxic microenvironments in inflamed tissue (Dvornikova *et al*, 2023; Solanki & Shah, 2024; Villareal & Xue, 2024). Intestinal ischemia-reperfusion injury similarly causes local oxygen deprivation, followed by its abrupt restoration upon reperfusion (Archontakis-Barakakis *et al*, 2025; Goncharov & Sharapov, 2023; Wu *et al*, 2018). Conditions such as COPD and obstructive sleep apnea (OSA) cause systemic hypoxia that also impairs oxygen delivery to the gut (May & Mehra, 2014; Wang *et al*, 2023), with OSA in particular producing repeated cycles of hypoxia and reoxygenation with each apneic episode. Fluctuations in oxygen availability are therefore a recurring feature of intestinal physiology in both health and disease. The mechanisms by which the intestinal epithelium maintains homeostasis in response to other environmental stresses, including nutrient fluctuation, enteric pathogens and ingested toxins, have been studied extensively (Beumer & Clevers, 2016; Hageman *et al*, 2020; Zwick *et al*., 2019). In contrast, how the epithelium responds to changing oxygen availability remains poorly understood.

The *Drosophila* midgut has emerged as a powerful and genetically tractable model for studying intestinal homeostasis and repair. As in the mammalian intestine, the adult fly midgut consists of a single-layered epithelium of large, absorptive enterocytes (ECs), which make up the bulk of the tissue, alongside a smaller population of secretory enteroendocrine cells (Takashima & Hartenstein, 2012; Wang & Hou, 2010). Both cell types arise from multipotent ISCs, which divide to produce transient progenitors that differentiate into replacement enterocytes or enteroendocrine cells (Guo & Ohlstein, 2015; Hou, 2010). ISC proliferative activity is dynamically scaled to the level of epithelial demand, ranging from slow homeostatic turnover that replaces the epithelium over the course of weeks, to rapid damage-induced regeneration in which ISC proliferation rates increase markedly and the entire epithelium can be reconstituted within days (Biteau *et al*., 2011; Colombani & Andersen, 2020; Jiang & Edgar, 2011; Li & Jasper, 2016; Xu *et al*, 2011; Zhang & Edgar, 2022).

Extensive work in this system has investigated how intestinal regeneration is triggered following epithelial damage. Much of this work has relied on enteric pathogens, chemical damaging agents, and genotoxins to define the mechanisms linking EC injury to ISC-mediated repair (Boumard & Bardin, 2021; Jiang *et al*, 2016; Zhang & Edgar, 2022; Zhou & Boutros, 2023). A central theme emerging from these studies is that damaged enterocytes actively promote their own replacement through the induction of the cytokines Unpaired 2 and Unpaired 3 (Upd2/3). These secreted ligands activate the conserved JAK-STAT signaling pathway in neighboring ISCs and progenitor cells, stimulating their proliferation and differentiation (Beebe *et al*, 2010; Buchon *et al*, 2009a; Buchon *et al*, 2009b; Jiang *et al*, 2009; Osman *et al*, 2012; Patel *et al*, 2019; Zhou *et al*, 2013). Upd production can be induced by a range of damage-responsive signaling pathways, including ROS, JNK (Biteau *et al*, 2008; Buchon *et al*., 2009a; Jiang *et al*., 2009; Mundorf *et al*, 2019; Patel *et al*, 2015), p38 (Patel *et al*., 2019; Santabarbara-Ruiz *et al*, 2015), and Hippo/Yorkie (Houtz *et al*, 2017; Li *et al*, 2014; Ren *et al*, 2010; Shaw *et al*, 2010; Staley & Irvine, 2010). In this way, damaged enterocytes act not only as targets of injury but also as the source of the regenerative signal, with the pathways that respond to the damage also driving cytokine induction.

Like all *Drosophila* tissues, the gut is supplied with oxygen directly by the tracheal system, a network of air-filled tubes whose terminal branches extend into the intestine and physically support its three-dimensional architecture within the abdomen (Blackie *et al*, 2024; Centanin *et al*, 2008). Following intestinal damage, the tracheal network remodels and branches toward the site of injury, and trachea-derived factors can also modulate the repair response (Li *et al*, 2013; Perochon *et al*, 2021; Tamamouna *et al*, 2021). These studies show how the tracheal network responds to intestinal damage, but leave open how the epithelium itself responds when oxygen availability changes.

Here, we use the adult Drosophila gut to investigate how the intestinal epithelium senses and responds to hypoxia-reoxygenation. We find that hypoxia activates AMPK-p53 signaling in a subset of enterocytes, priming them for death, while inducing Upd cytokine production in a largely separate population that activates STAT signaling in neighboring ISCs, priming them for proliferation. However, enterocyte death and ISC proliferation only occur upon reoxygenation. This couples the timing of cell loss and stem cell-mediated repair to the return of oxygen, rather than to the hypoxic insult itself. This response is also independent of hypoxia-inducible factor 1α (HIF-1α) in enterocytes but requires HIF-1α in progenitor cells for their reoxygenation-triggered proliferation.

## Results

### Acute hypoxia-reoxygenation drives reversible intestinal resizing

To investigate the effects of acute hypoxia on intestinal morphology, we exposed adult female flies to sublethal durations of hypoxia (1% oxygen) and compared midgut length to normoxic controls. We found that hypoxia exposure led to a rapid decrease in gut length, with a ∼30% reduction within 2 hours of exposure that persisted at 6 and 16 hours of continuous hypoxia (Fig. 1A–B). To examine whether this effect was reversible, we exposed flies to 6 hours of hypoxia and monitored gut length upon return to ambient oxygen. We found that reoxygenation led to a gradual recovery of gut length to normoxic values by 24 hours, which was maintained through 48 hours of reoxygenation (Fig. 1C). Together these results show that acute hypoxia drives a rapid reduction in intestinal size that is fully reversible upon reoxygenation.

**Figure 1:**
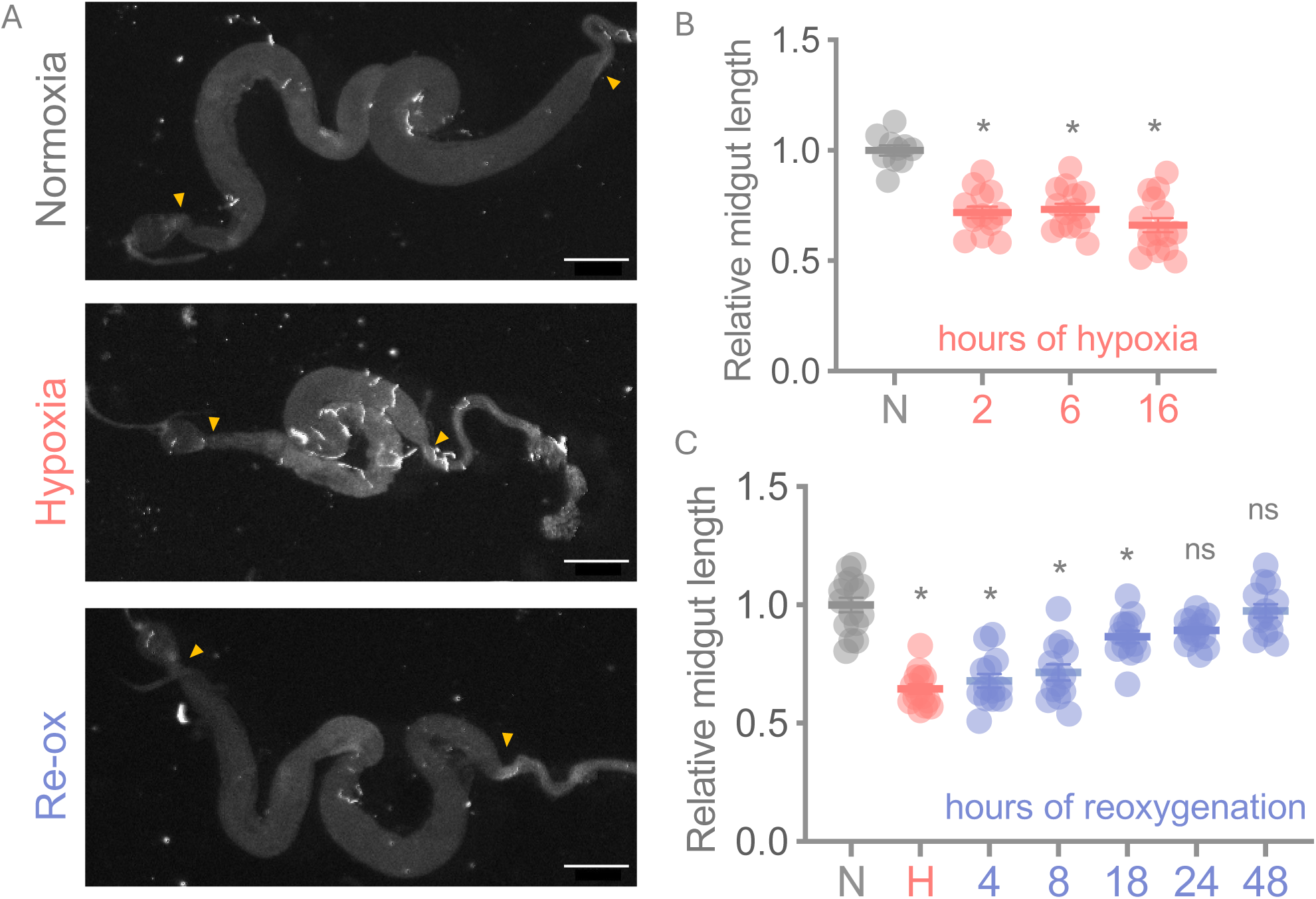
Acute hypoxia-reoxygenation leads to rapid intestinal resizing. (A, B) Age-matched flies (*w^1118^*) were maintained in normoxia or hypoxia (1% oxygen) and dissected for gut length measurements in the region flanked by yellow arrowheads. Brightfield images (A) were quantified for midgut length using Fiji and relative midgut length is plotted in (B). Data represents mean ± SEM, n =10-15 independent guts for each experimental condition. Scale bar = 1000 *μ*m. *p<0.05 unpaired t-test. (A, C) To test whether the hypoxia effect on gut length was reversible, age-matched flies (*w^1118^*) were maintained either at normoxia or exposed to acute hypoxia (1% oxygen) for 6 hours and then reoxygenated. Brightfield images were captured for guts dissected at various timepoints to measure gut length in the region flanked by yellow arrowheads. As quantified in (C), gut length recovered by 24 hours of reoxygenation following hypoxia exposure. Each data point depicts individual tissues and data represents mean ± SEM, n = 10-15 guts per experimental condition. One-way ANOVA *p<0.05 compared to normoxic controls. ns: not significant.

### Caspase activation in hypoxia primes enterocytes for apoptosis upon reoxygenation

Given the rapid hypoxia-induced change in intestinal size, we next asked what happens to individual cells of the intestinal epithelium during hypoxia and reoxygenation. Since hypoxia is a potent cellular stressor (Majmundar *et al*, 2010; Zhou & Haddad, 2013), we first asked whether it causes cellular damage by staining for the Death caspase-1, Dcp-1 (McCall & Steller, 1998; Song *et al*, 1997). This revealed a marked increase in caspase-positive cells in hypoxic intestines compared to normoxic controls, with caspase activity confined to large polyploid ECs (Fig. 2A).

**Figure 2:**
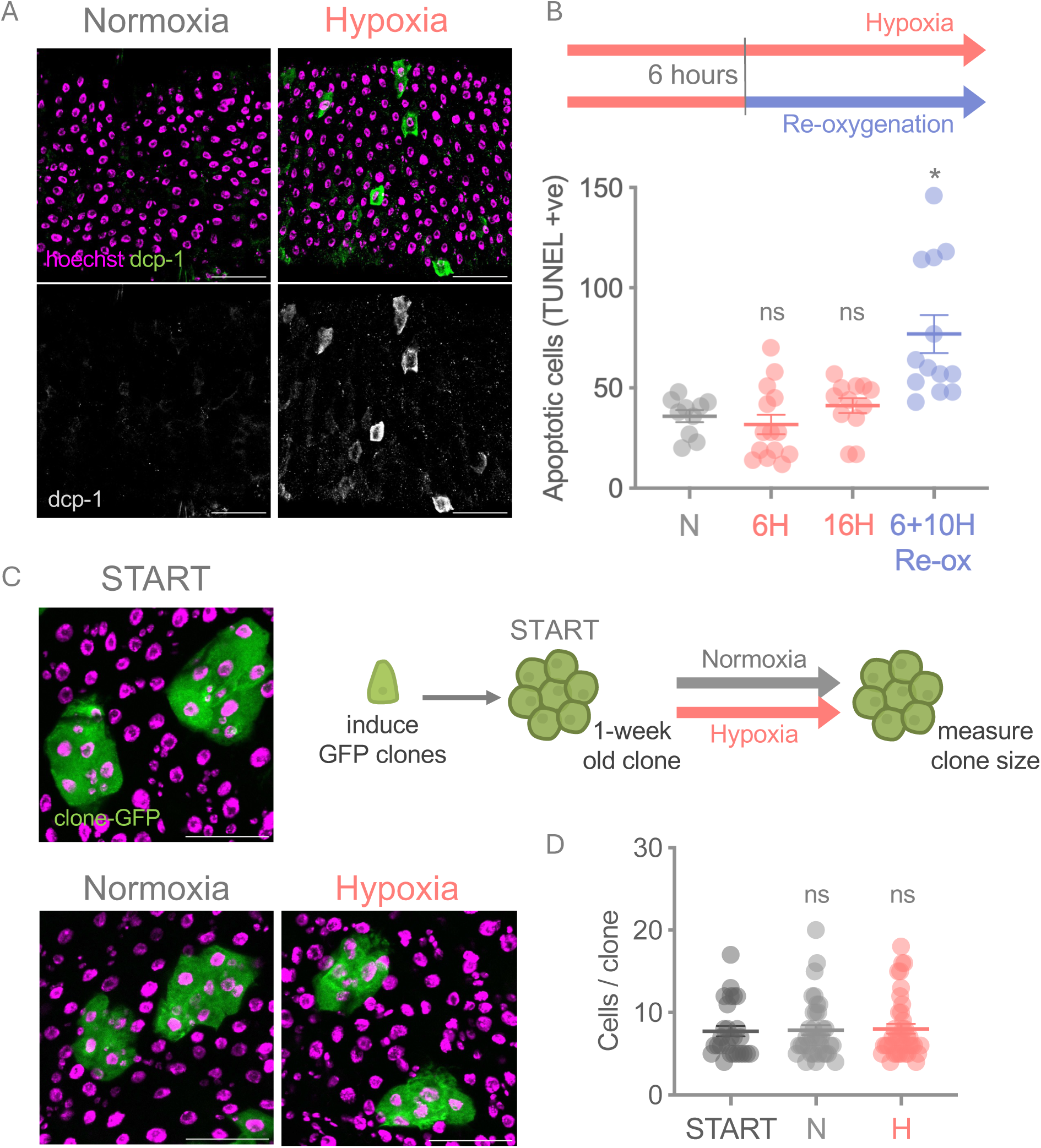
Acute hypoxia upregulates caspase activity, but reoxygenation is required for cell apoptosis. **(A)** Guts were stained for Dcp-1 (Death caspase-1) to measure caspase activity in normoxic and hypoxic intestines. Hypoxia induced caspase activity in cells of the intestinal epithelium. Scale bar = 50 µm. (B) Schematic of the hypoxia and reoxygenation regimens, with TUNEL labeling of fragmented DNA quantified below. Continuous hypoxia for 16 hours did not increase apoptotic cell numbers compared to normoxia, whereas 6 hours of hypoxia followed by 10 hours of reoxygenation did, indicating that apoptosis requires reoxygenation. Data represents individual tissues showing mean ± SEM, n = 10-15 guts per experimental condition. One-way ANOVA *p<0.05, ns: not significant. (C-D) Clonal analysis was used to measure cell loss from the epithelium during the hypoxic period. Flies were heat-shocked to induce GFP clones which were allowed to grow for a week (6-10 cells/clone). Clone size was then measured prior to hypoxia exposure (START) and at the end of 6 hours of hypoxia (1% oxygen) or normoxia as seen in the representative images (C). Clone size remained unchanged post hypoxia exposure as quantified in (D). Data represents mean ± SEM, n = 25-40 clones analyzed from 10-13 independent guts for each experimental condition. Scale bar = 50 *μ*m. ns: not significant, unpaired t-test.

To ask whether this caspase induction was accompanied by active cell death, we stained for TUNEL, a marker of late-stage apoptosis. TUNEL-positive cell numbers were not elevated at the end of 6 hours of hypoxia compared to normoxic controls (Fig. 2B). Caspase activation was therefore not accompanied by cell death during hypoxia. Instead, we saw that TUNEL-positive cells accumulated specifically during the reoxygenation phase when we returned flies to ambient oxygen following hypoxia exposure. To test whether this delayed induction of apoptosis was simply a consequence of the time required to initiate and execute apoptosis, we compared flies maintained under continuous hypoxia for 16 hours to flies subjected to 6 hours of hypoxia followed by 10 hours of reoxygenation. Only the reoxygenated group showed elevated TUNEL labeling (Fig. 2B), demonstrating that reoxygenation is required for cell death to occur.

To further examine whether cells were being lost from the epithelium during hypoxia, we examined the fate of GFP-marked cell clones. We used the MARCM (Mosaic Analysis with a Repressible Cell Marker) system to induce stem cell-derived clones throughout the intestine (Wu & Luo, 2006), allowed these to grow for one week to generate clones of 6-10 cells, and then exposed flies to normoxia or hypoxia for 6 hours. The number of cells per clone was not different between normoxic and hypoxic conditions (Fig. 2C–D), indicating that hypoxia does not cause detectable cell loss over this period.

Since cells were not lost from the epithelium during hypoxia, the rapid reduction in gut length must have another basis. We therefore examined enterocyte morphology. We found that hypoxia caused enterocytes to become narrower and taller, with basal cell area reduced by approximately 40% and apical-basal cell length increased by approximately 20–30% relative to normoxic controls (Fig. S1). These changes were evident by 2 hours and maintained at 6 hours, matching the time course of the reduction in gut length.

Together these results show that hypoxia induces caspase activation without cell loss, priming enterocytes for death upon reoxygenation, and indicate that the rapid reduction in gut length instead reflects a change in enterocyte shape.

### Enterocyte apoptosis is required for the reoxygenation-induced burst of ISC proliferation

We next examined the recovery of intestinal length upon reoxygenation. Because enterocyte apoptosis occurred during reoxygenation, we reasoned that new cells must be produced to compensate for this loss and to account for the recovery in gut length. To examine this, we used the MARCM system to generate single-cell ISC clones immediately prior to hypoxia exposure and measured clone size after 6 hours of hypoxia followed by 18 hours of reoxygenation, comparing to clones from flies maintained under normoxia for the same 24-hour duration. We found that hypoxia-reoxygenation led to a significant increase in the number of cells per clone compared to normoxia (Fig. 3A–B), indicating increased stem cell divisions during reoxygenation. To directly quantify ISC proliferative activity, we measured mitotic stem cells by phospho-histone H3 (pH3) staining across the hypoxia and reoxygenation time course. ISC divisions were reduced below normoxic levels during the 6 hour hypoxia exposure, but increased upon reoxygenation, peaking at 18 hours before returning toward baseline by 48 hours (Fig. 3C).

**Figure 3:**
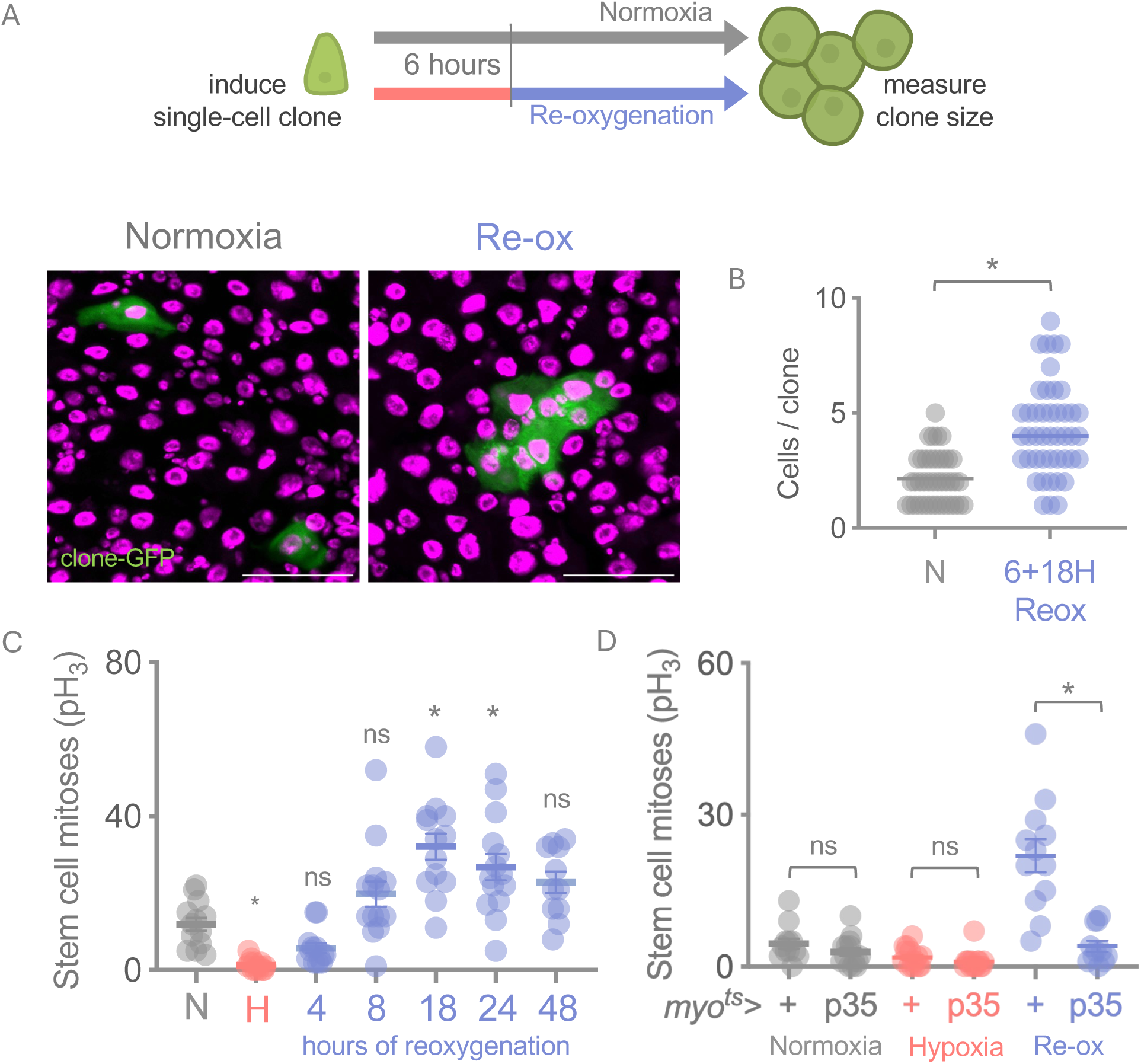
Cell loss is coupled to cell gain during reoxygenation. (A-B) Clonal analysis was used to measure cell turnover in the epithelium during the reoxygenation period. Flies were heat-shocked to induce single cell GFP clones immediately before hypoxia exposure. Clone size was measured after exposing flies to 6 hours of hypoxia, followed by 18 hours of reoxygenation, and compared to control clone size from flies maintained at ambient oxygen for 24 hours. Data represents mean ± SEM, n = 25-40 clones analyzed from 10-13 independent guts for each experimental condition. Scale bar = 50 *μ*m. *p<0.05 unpaired t-test. (C) Guts were stained with phospho-histone H3 (pH3) mitosis marker to measure proliferating stem cells in the epithelium at various timepoints. Hypoxia exposure suppressed baseline stem cell divisions. Upon reoxygenation, stem cell divisions were elevated, peaking at 18 hours after hypoxia exposure. Each point depicts total proliferating stem cells in individual tissues and data represents mean ± SEM, n = 10-15 guts per experimental condition. One-way ANOVA *p<0.05, ns: not significant compared to normoxia. (D) Adult-enterocyte specific expression of baculoviral gene p35 did not affect stem cell division in normoxia or hypoxia, compared to controls. Reoxygenation-associated proliferation was blunted upon enterocyte-specific expression of p35, compared to controls. Each point depicts total proliferating stem cells in individual tissues and data represents mean ± SEM, n = 10-15 guts per experimental condition. *p<0.05, ns: not significant, two-way ANOVA with Tukey’s multiple comparisons test.

Since enterocyte apoptosis and ISC proliferation both occur during reoxygenation, we asked whether these events are coupled. We expressed the baculoviral caspase inhibitor p35 (HAY *et al*, 1994) in adult enterocytes using the temperature-sensitive driver *myo1A-Gal4^ts^* (*myo^ts^>p35*) to block apoptosis, and measured ISC mitoses at 18 hours of reoxygenation following hypoxia exposure. We found that p35-mediated blockade of enterocyte apoptosis had no effect on ISC divisions during normoxia or hypoxia, but reduced the proliferative surge during reoxygenation to near-baseline levels (Fig. 3D).

Together these results show that ISC divisions are suppressed during hypoxia and increase upon reoxygenation, coinciding with the phase in which enterocyte apoptosis occurs. Blocking enterocyte apoptosis reduced ISC divisions to near-baseline levels, showing that enterocyte loss and stem cell-mediated replacement are coupled during reoxygenation.

### An AMPK-p53-*reaper* pathway mediates enterocyte apoptotic priming in hypoxia

We next investigated the signaling mechanisms that link hypoxia to caspase induction in enterocytes. We first examined a role for HIF-1α, the conserved central mediator of adaptive responses to hypoxia, which is degraded in the presence of oxygen and stabilized when oxygen becomes limiting (Lavista-Llanos *et al*, 2002). In the mammalian intestine, HIF-1α signaling supports barrier function, epithelial metabolism and the response to inflammation (Brown & Taylor, 2018; Taylor & Colgan, 2017). We used RNAi to knock down expression of *sima* (the *Drosophila* homolog of HIF-1α) specifically in enterocytes using the mex-Gal4 driver and a UAS-*sima* RNAi line that we previously showed induces strong knockdown of *sima* target genes (Ding *et al*, 2026). Hypoxia-induced caspase activity appeared unchanged following sima knockdown (Fig. S2A), and reoxygenation-induced stem cell proliferation was unaffected (Fig. S2B), arguing against a role for HIF-1α in the enterocyte effects we see upon hypoxia-reoxygenation.

AMPK is a conserved energy-sensing kinase activated by increases in the cellular AMP and ADP to ATP ratio (Hardie *et al*, 2012; Pan & Hardie, 2002) and has been shown to mediate many of the cellular effects of hypoxia (Dengler, 2020; Liu *et al*, 2006). We therefore examined a potential role for enterocyte AMPK in the effects of hypoxia-reoxygenation on intestinal biology. To do so, we used the mex-Gal4^ts^ driver to express a kinase-dead dominant-negative AMPK (*mex^ts^>AMPK^DN^*) to inhibit AMPK signaling in enterocytes (Johnson *et al*, 2010). These flies exhibited a significant reduction in caspase-positive cells during hypoxia compared to controls (*mex^ts^*>+), indicating that AMPK signaling is required for the hypoxia-induced caspase activation in enterocytes (Fig. 4A–B). Consistent with the coupling between enterocyte apoptosis and ISC proliferation established above, AMPK^DN^ expression also blunted the reoxygenation-driven induction of stem cell divisions (Fig. 4C). Hypoxia-induced shortening was unaffected in these flies, but recovery of intestinal length was impaired upon reoxygenation (Fig. 4D).

**Figure 4:**
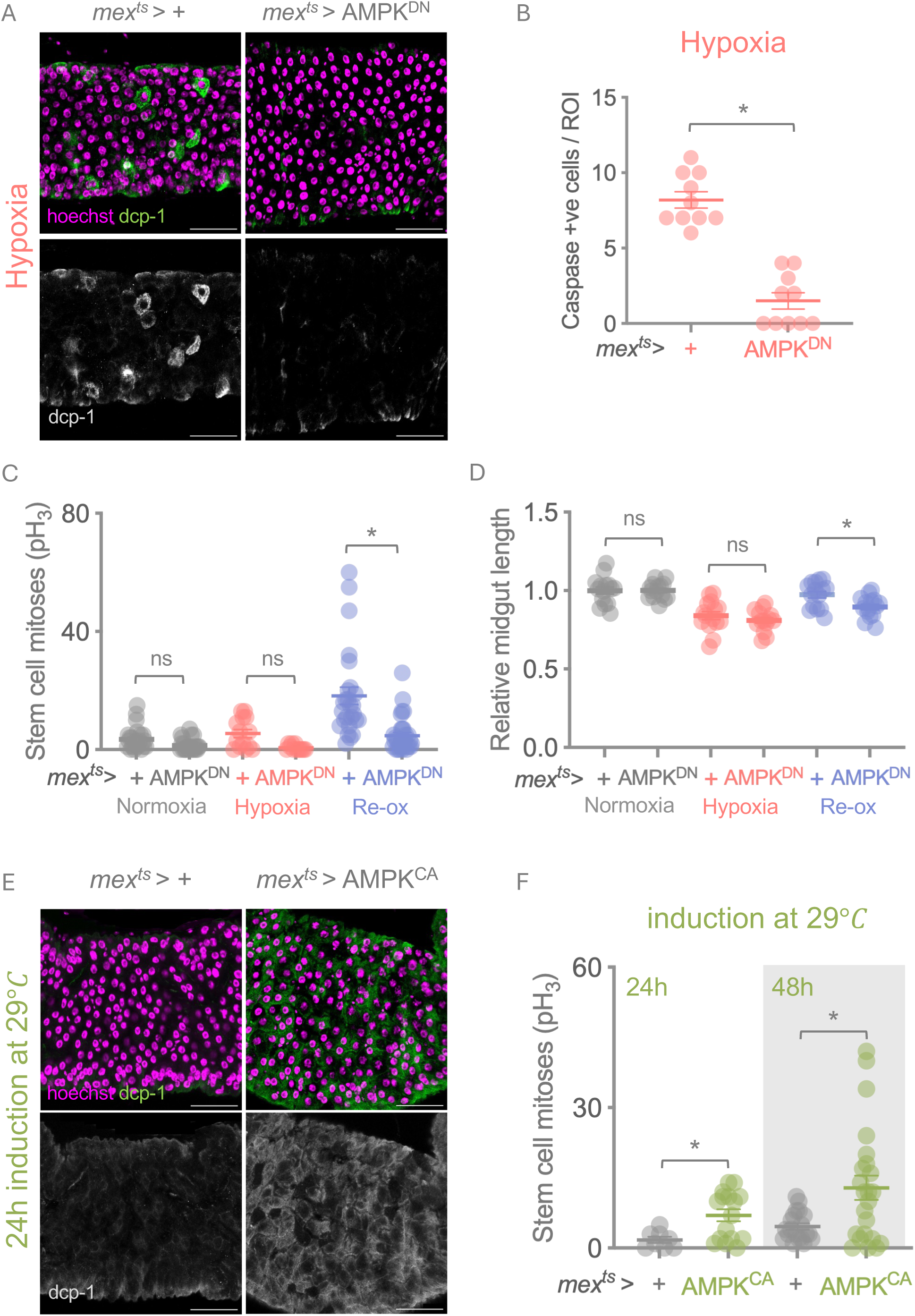
AMPK coordinates caspase induction in hypoxia. (A-B) Guts were stained with Dcp-1 to measure caspase activity in intestines exposed to hypoxia. Control intestines induce caspase activity in large polyploid cells. Overexpression of a kinase dead AMPK (*AMPK^DN^*) in adult enterocytes resulted in suppression of caspase induction during the hypoxic period, as quantified in (B) *p<0.05 (unpaired t-test). (C-D) Adult enterocyte expression of *AMPK^DN^* resulted in suppressed reoxygenation-associated stem cell divisions, and impaired gut length recovery in reoxygenation, compared to controls. Each point depicts total proliferating stem cells, or gut lengths, in individual tissues and data represents mean ± SEM, n = 10-15 guts per experimental condition. *p<0.05, ns: not significant (two-way ANOVA with Tukey’s multiple comparisons test). (E-F) A constitutive active version of AMPKα (*AMPK^CA^*) was expressed in adult enterocytes for 24 hours. This was sufficient to induce caspase activity, as visualized by intestines stained for Dcp-1. Induction of enterocyte *AMPK^CA^* expression also led to an increase in the levels of stem cell proliferation as quantified in (F). This suggests that AMPK activity is sufficient to induce caspase activity and downstream increase stem cell proliferation. Each data point depicts total proliferating stem cells in individual tissues and data represents mean ± SEM, n = 10-15 guts per experimental condition. Scale bar = 50 *μ*m. *p<0.05 (unpaired t-test).

To determine whether AMPK activity is sufficient to induce caspase activation, we used *mex-Gal4^ts^* to express a constitutively active AMPKα (*mex^ts^>AMPK^CA^*) (Swick *et al*, 2013) in adult enterocytes under normoxic conditions. Expression in the absence of hypoxia was sufficient to elevate cleaved caspase signal at 24 hours (Fig. 4E) and to increase mitotic stem cell numbers at both 24 and 48 hours (Fig. 4F). Enterocyte AMPK activity is therefore sufficient to drive both caspase activation and ISC proliferation when oxygen is present.

Having established AMPK as an upstream regulator of caspase induction in enterocytes, we next sought to identify how AMPK promotes pro-apoptotic signaling. p53 induces expression of pro-apoptotic genes to drive apoptosis (Brodsky *et al*, 2000; Dichtel-Danjoy *et al*, 2013; Mollereau & Ma, 2014), and AMPK has been reported to function through p53 in the context of metabolic stress (Ingaramo *et al*, 2020). We therefore examined whether p53 mediates AMPK-dependent caspase induction in hypoxia. *p53* mRNA levels were strongly elevated in hypoxic intestines relative to normoxic controls, and enterocyte knockdown of AMPK (*mex^ts^*>*AMPK* RNAi) significantly reduced this induction (Fig. 5A). Enterocyte-specific knockdown of p53 using the constitutive *mex*-Gal4 driver (*mex*>*p53* RNAi) reduced the number of caspase-positive cells in hypoxia compared to controls (*mex*>+), phenocopying the effects of AMPK^DN^ expression (Fig. 5B). In *Drosophila*, the pro-apoptotic gene *reaper* is a transcriptional target of p53 that drives caspase activation (Brodsky *et al*., 2000; White *et al*, 1994; White *et al*, 1996). Both AMPK^DN^ expression and *p53* knockdown significantly suppressed hypoxia-induced *reaper* transcription (Fig. 5C–D). Finally, *p53* knockdown in enterocytes blunted ISC proliferation during reoxygenation (Fig. 5E), mirroring the effects of AMPK^DN^ and p35.

**Figure 5:**
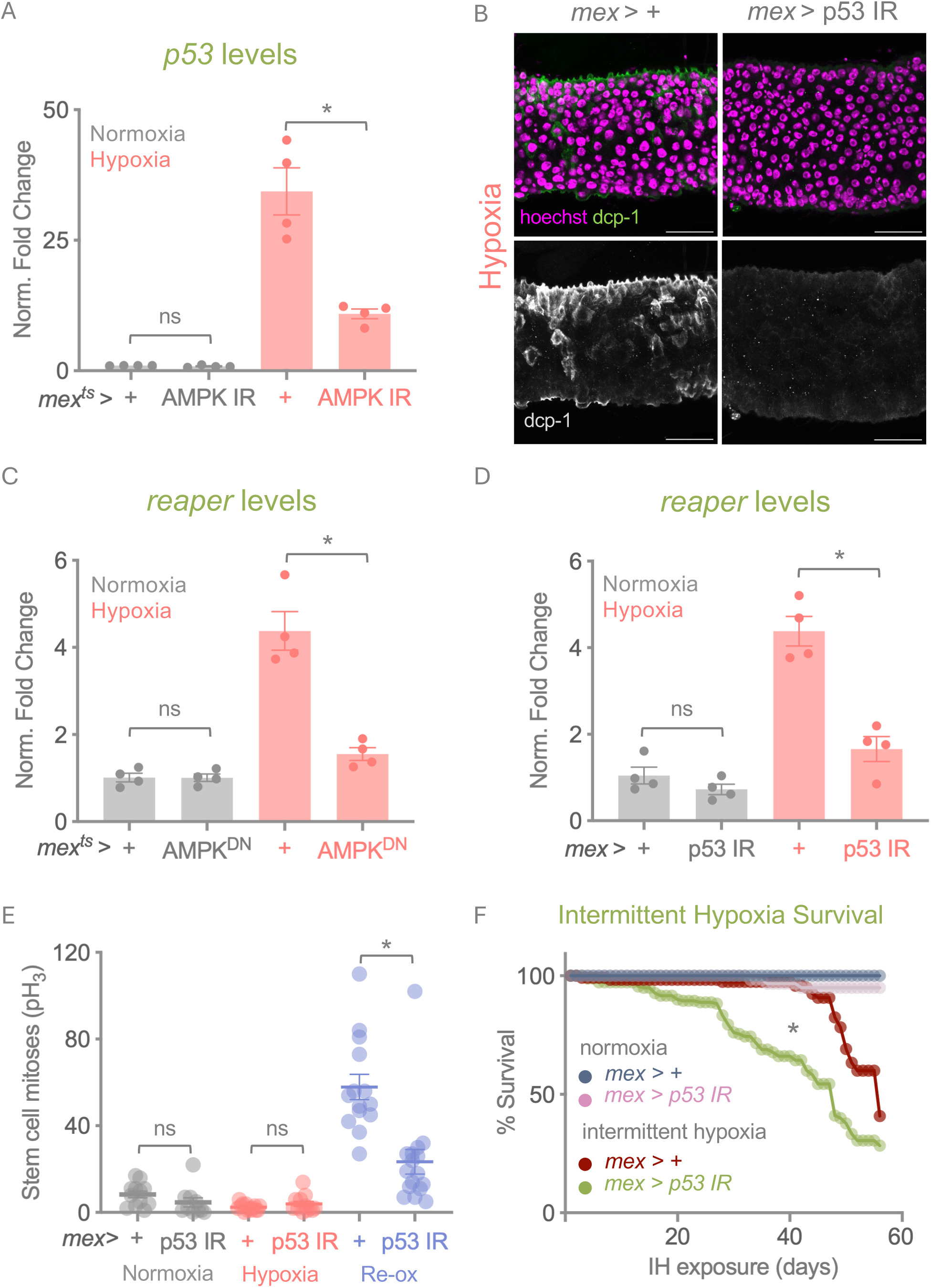
AMPK mediates pro-apoptotic signaling in hypoxia through p53. (A) Guts from flies exposed to 6 hours of acute hypoxia were dissected for qPCR analysis. *p53* is transcriptionally upregulated in control hypoxic guts. Adult-enterocyte-specific expression of dsRNA to *AMPK* resulted in suppression of *p53* transcript levels in hypoxia. Data represents mean ± SEM, N=4 independent groups of samples (5 animals per group) per experimental condition. *p<0.05, ns: not significant (two-way ANOVA with Tukey’s multiple comparisons test). Data points represent independent samples. (B) Guts from flies exposed to 6 hours of acute hypoxia were dissected and stained for Dcp-1. Enterocyte-specific expression of dsRNA to *p53* resulted in suppression of caspase induction in hypoxic intestines. Scale bar = 50 *μ*m. (C-D) AMPK mediates caspase induction in hypoxia through upregulating pro-apoptotic gene *reaper* transcript levels. *reaper* transcripts are induced in hypoxia. Enterocyte-specific expression of (C) *AMPK^DN^*, or (D) dsRNA to *p53*, results in suppression of *reaper* transcript in hypoxia. Data represents mean ± SEM, N=4 independent groups of samples (5 animals per group) per experimental condition. *p<0.05, ns: not significant (two-way ANOVA with Tukey’s multiple comparisons test). Data points represent independent samples. (E) Enterocyte-specific knockdown of p53 blunts stem cell proliferation during reoxygenation compared to controls. Each data point depicts total proliferating stem cells in individual tissues and data represents mean ± SEM, n = 10-15 guts per experimental condition. *p<0.05, ns: not significant (two-way ANOVA with Tukey’s multiple comparisons test). (F) Survival curves for flies exposed to intermittent hypoxia (6 hours of 1% Oxygen, three times a week). Enterocyte-specific knockdown of p53 had no effect on survival in normoxia. Exposure to intermittent hypoxia reduced lifespan in controls, which was further exacerbated in enterocyte-specific p53 knockdown. n = 40-50 in normoxia. n = 140-180 in intermittent hypoxia. *p<0.05 (Mantel-Cox log rank test).

To examine the organismal consequence of disrupting this response, we knocked down p53 in enterocytes and examined viability following repeated cycles of hypoxia-reoxygenation. Flies were exposed to repeated bouts of 6 hours of hypoxia, three times per week, with return to normoxia between bouts, to test the consequences of repeated hypoxia-reoxygenation cycles of the kind that occur in human pathologies such as obstructive sleep apnea. Control flies (*mex*>+) exposed to intermittent hypoxia showed reduced lifespan compared to flies maintained in normoxia, and this effect was markedly exacerbated in flies with enterocyte-specific *p53* knockdown (Fig. 5F).

Together these results define an AMPK-p53-*reaper* signaling axis linking hypoxia to enterocyte apoptotic priming, with AMPK both necessary and sufficient for caspase induction and acting upstream of p53 and *reaper*. Disrupting this axis blunted ISC proliferation upon reoxygenation and reduced survival under intermittent hypoxia.

### Hypoxia primes stem cells for proliferation through progenitor HIF-1α and enterocyte-derived Upd signaling

We next asked how the stem cell compartment is prepared for the proliferative burst that occurs upon reoxygenation. Our earlier results indicated that HIF-1α was not required in enterocytes for caspase activation during hypoxia or for the consequent stem cell proliferation during reoxygenation. However, qPCR analysis of isolated intestines showed that hypoxia increased expression of known *sima* target genes (Centanin *et al*., 2008; Lavista-Llanos *et al*., 2002), including *branchless* (*bnl*) and *Lactate dehydrogenase* (*Ldh*) (Fig. S2C, D), indicating that *sima* activity is induced within the intestine. To explore the spatial features of this induction, we examined a reporter line expressing endogenously tagged Ldh (*ldh::gfp*). Ldh-GFP was most pronounced in small cells with the morphology of progenitors (Fig. 6A), and the proportion of progenitors positive for Ldh-GFP increased in hypoxia and at 2 hours of reoxygenation (Fig. 6B). Induction of this *sima* target therefore occurs largely in progenitor-like cells rather than in enterocytes. To examine the consequence of this induction, we used the temperature-sensitive progenitor-specific driver *esg-Gal4^ts^* to knock down *sima* (*esg^ts^*>*sima* RNAi). Progenitor-specific *sima* knockdown had no effect on stem cell proliferation in normoxia but significantly decreased the burst of stem cell proliferation induced by reoxygenation (Fig. 6C), indicating a progenitor-autonomous requirement for HIF-1α in the reoxygenation-induced proliferative response.

**Figure 6:**
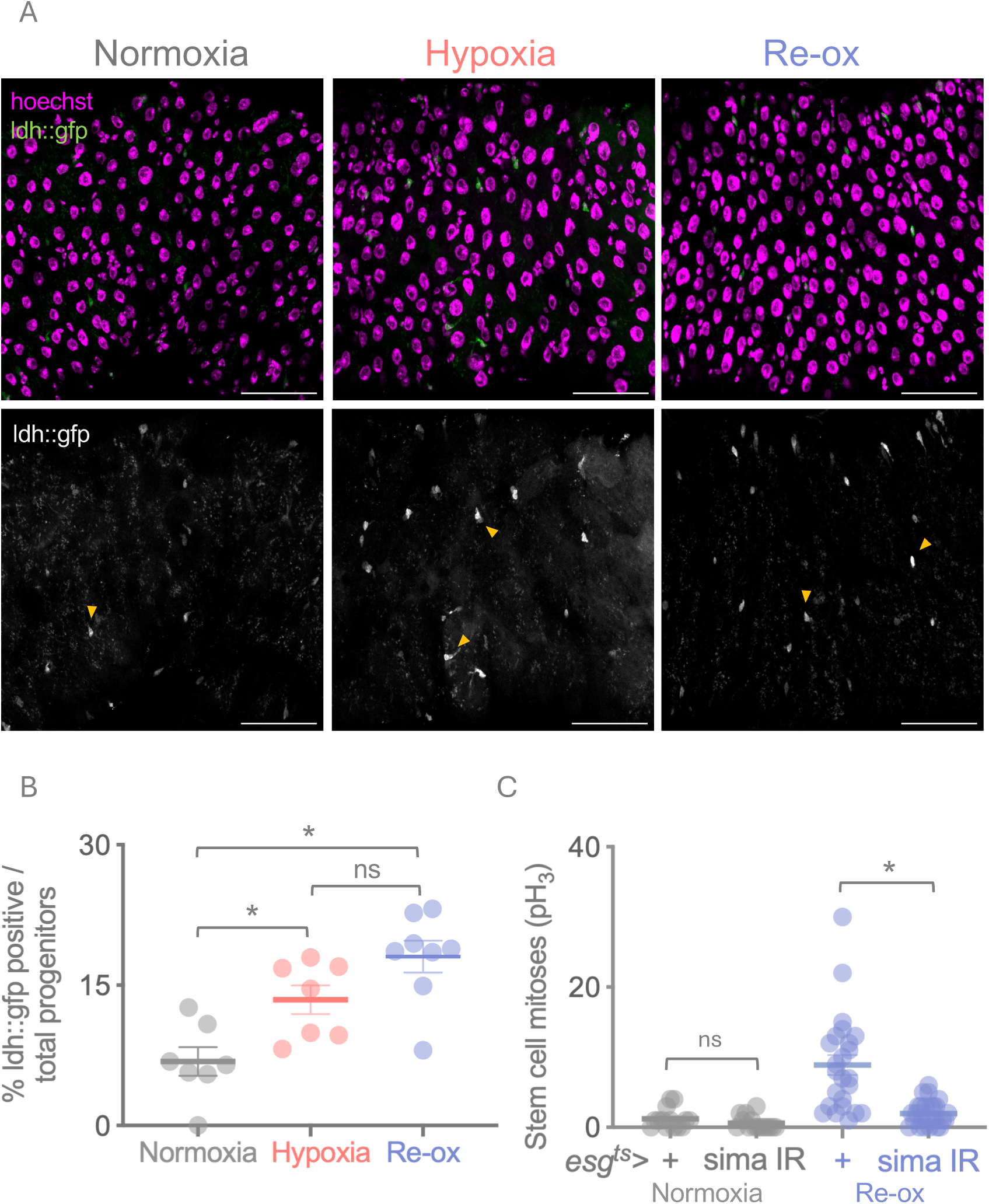
Progenitor cell HIF-1α is required for stem cell divisions during reoxygenation. (A) Lactate dehydrogenase (Ldh) expression (*ldh::gfp)* is largely localized to intestinal progenitor cells. Yellow arrows indicate cells considered positive for Ldh expression as quantified in (B). Scale bar = 50 *μ*m. Data represents mean ± SEM, quantified over 7-8 individual tissues. *p<0.05, ns: not significant (One-way ANOVA). (C) Expression of dsRNA to HIF-1α in adult intestinal progenitor cells had no impact on stem cell divisions during normoxia or hypoxia, but resulted in suppressed stem cell divisions during reoxygenation. Each data point depicts total proliferating stem cells in individual tissues and data represents mean ± SEM, n = 10-15 guts per experimental condition. *p<0.05, ns: not significant (two-way ANOVA with Tukey’s multiple comparisons test).

We also found that mRNA levels of the cytokines *upd2* and *upd3* were strongly upregulated in hypoxic intestines relative to normoxic controls (Fig. 7A). When we examined expression of an *upd3* reporter (*upd3*-*Gal4*>*UAS-GFP*), we saw elevated GFP expression localized predominantly to the large, polyploid enterocytes (Fig. 7B). This matches our previous finding that hypoxic induction of *upd3* in the gut occurs predominantly in enterocytes (Ding *et al*., 2026). Using a STAT-GFP transcriptional reporter as a read-out of Upd-JAK-STAT signaling (Bach *et al*, 2007), we found elevated STAT activity in the small progenitor cells during hypoxia exposure (Fig. 7C–D). These results indicate that in low oxygen a population of enterocytes produces Upd cytokines that stimulate JAK-STAT activity in gut progenitor cells. JAK-STAT activation typically stimulates ISC proliferation, yet ISC mitoses were suppressed during this period. We therefore considered that JAK-STAT upregulation in ISCs during hypoxia primes them for proliferation upon reoxygenation. To test this, we measured ISC divisions and intestinal length during reoxygenation in whole-body *upd2, upd3* double mutants (*upd2,3*Δ) (Osman *et al*., 2012). Loss of Upd signaling had no effect on ISC proliferation in normoxia but blunted the reoxygenation-associated wave of ISC divisions and prevented intestinal length recovery (Fig. 7E–F). Knockdown of *Stat92E* with the temperature-sensitive progenitor-specific driver esg-Gal4^ts^ (*esg^ts^*>*Stat92E* RNAi) phenocopied the *upd2,3*Δ defects, with suppressed ISC proliferation and impaired intestinal length recovery during reoxygenation (Fig. 7G–H). Because this manipulation was restricted to progenitor cells, intestinal length recovery during reoxygenation requires Upd-STAT signaling within the progenitor compartment.

**Figure 7:**
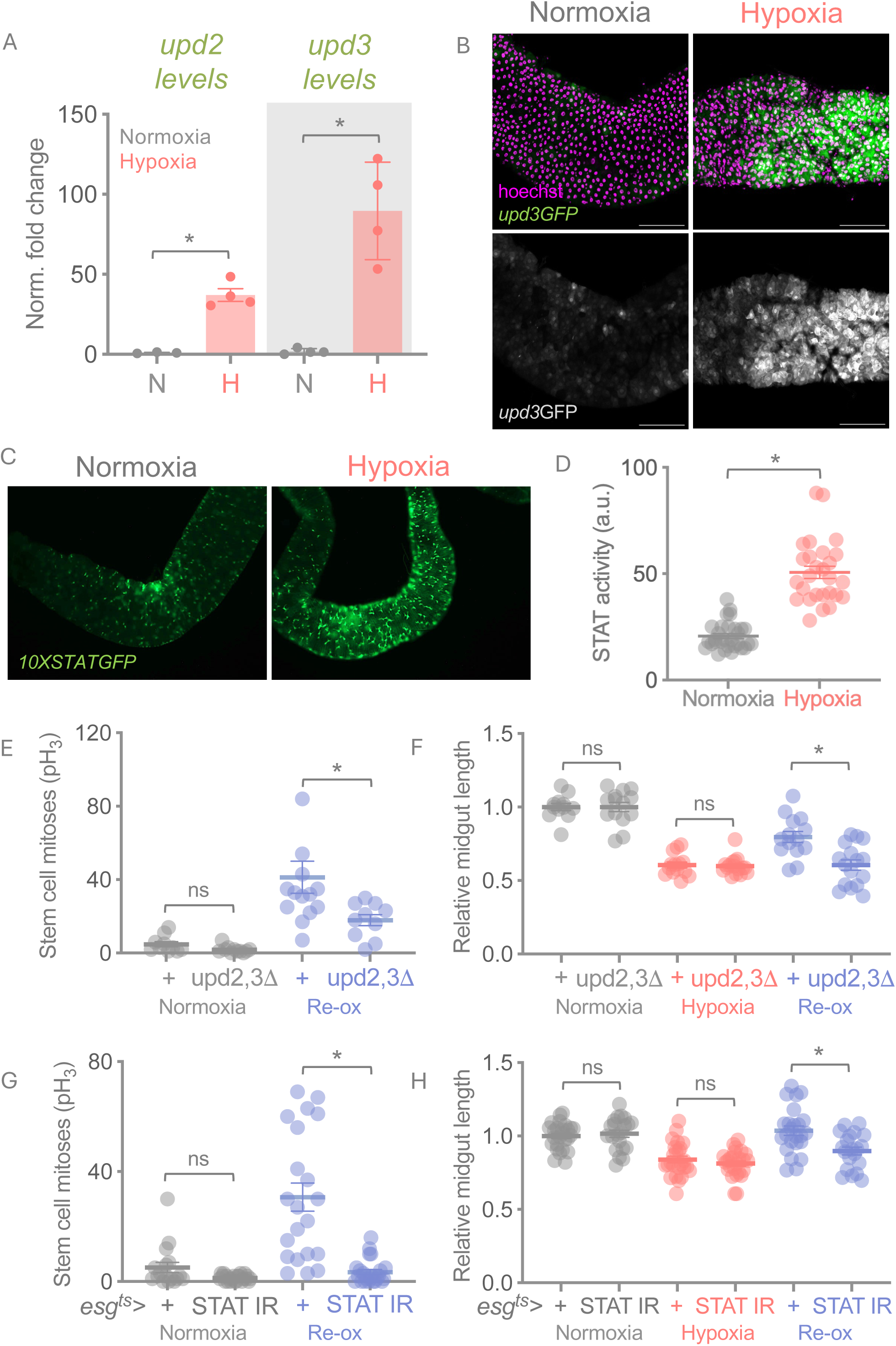
*Upd* cytokines upregulated in hypoxia are required for stem cell divisions in reoxygenation. (A) *unpaired 2 (upd2)* and *unpaired 3 (upd3)* transcripts are strongly upregulated in hypoxic intestines. Data represents mean ± SEM, N=4 independent groups of samples (5 animals per group) per experimental condition. *p<0.05 (unpaired t-test). Data points represent independent samples. (B) Representative images for *upd3-GFP* reporter activity show increased GFP fluorescence in hypoxia, primarily in large, polyploid enterocytes. Scale bar = 100 *μ*m. (C) Representative images for STAT92E activity (*10XSTAT GFP*) shows increased GFP fluorescence in hypoxia, primarily in smaller progenitor cells. Increase in STAT reporter activity in hypoxia is quantified in (D). *upd3* and *STAT92E* reporter activity suggest that Upd/JAK/STAT signaling is active in hypoxia. a.u. arbitrary units. *p<0.05 (unpaired t-test). (E-H) Upd/JAK/STAT signaling is required for stem cell mediated regeneration response in reoxygenation. Whole body mutants for *upd2,3* show blunted stem cell divisions (E), and reduced ability to recover intestinal length (F) in the same reoxygenation time frame as the controls. Similarly, progenitor-specific knockdown of *STAT92E* phenocopies whole body *upd2,3* mutants, showing strongly reduced stem cell divisions and impaired recovery of intestinal length. Each point depicts total proliferating stem cells, or gut lengths, in individual tissues and data represents mean ± SEM, n = 10-15 guts per experimental condition. *p<0.05, ns: not significant (two-way ANOVA with Tukey’s multiple comparisons test).

Hypoxia therefore activates HIF signaling within intestinal progenitor cells and induces enterocyte-derived Upd signaling that stimulates JAK-STAT activity in the same compartment. Although ISC proliferation is suppressed during hypoxia, both pathways are required for the wave of stem cell divisions upon reoxygenation, indicating that hypoxia initiates a second response that primes the intestine for repair.

### Apoptotic priming and Upd induction occur largely in separate enterocyte populations

Our results show that hypoxia triggers two responses in enterocytes. An AMPK-p53 stress response induces caspases and primes enterocytes for apoptosis, and an Upd response primes ISCs for proliferation. Neither is executed until oxygen is restored. Prevailing models of *Drosophila* intestinal repair propose that damaged enterocytes induce Upd cytokines to promote their own replacement (Houtz *et al*., 2017; Jiang *et al*., 2009; Mundorf *et al*., 2019; Patel *et al*., 2019; Santabarbara-Ruiz *et al*., 2015; Staley & Irvine, 2010; Zhang & Edgar, 2022; Zhou *et al*., 2013). We therefore asked whether the AMPK-p53-caspase pathway functions upstream of Upd induction, or whether these responses represent distinct branches of the hypoxic response.

We blocked this pathway at three levels in enterocytes, using *mex^ts^>AMPK* RNAi to deplete AMPK, *mex>p53* RNAi to deplete p53, and *mex^ts^>p35* to block caspase activity directly. None of these manipulations suppressed *upd2* or *upd3* transcript levels in hypoxia, and in some cases transcript levels were further elevated (Fig. 8A–C), showing that Upd cytokine induction is independent of the pathway that drives enterocyte caspase activation and apoptosis. To examine this further, we co-visualized cleaved Dcp-1 and *upd3* reporter expression at 2, 4 and 6 hours of hypoxia. Caspase-positive and *upd3*-expressing cells were largely non-overlapping at all three timepoints, despite both populations being confined to the large, polyploid enterocytes (Fig. 8D).

**Figure 8:**
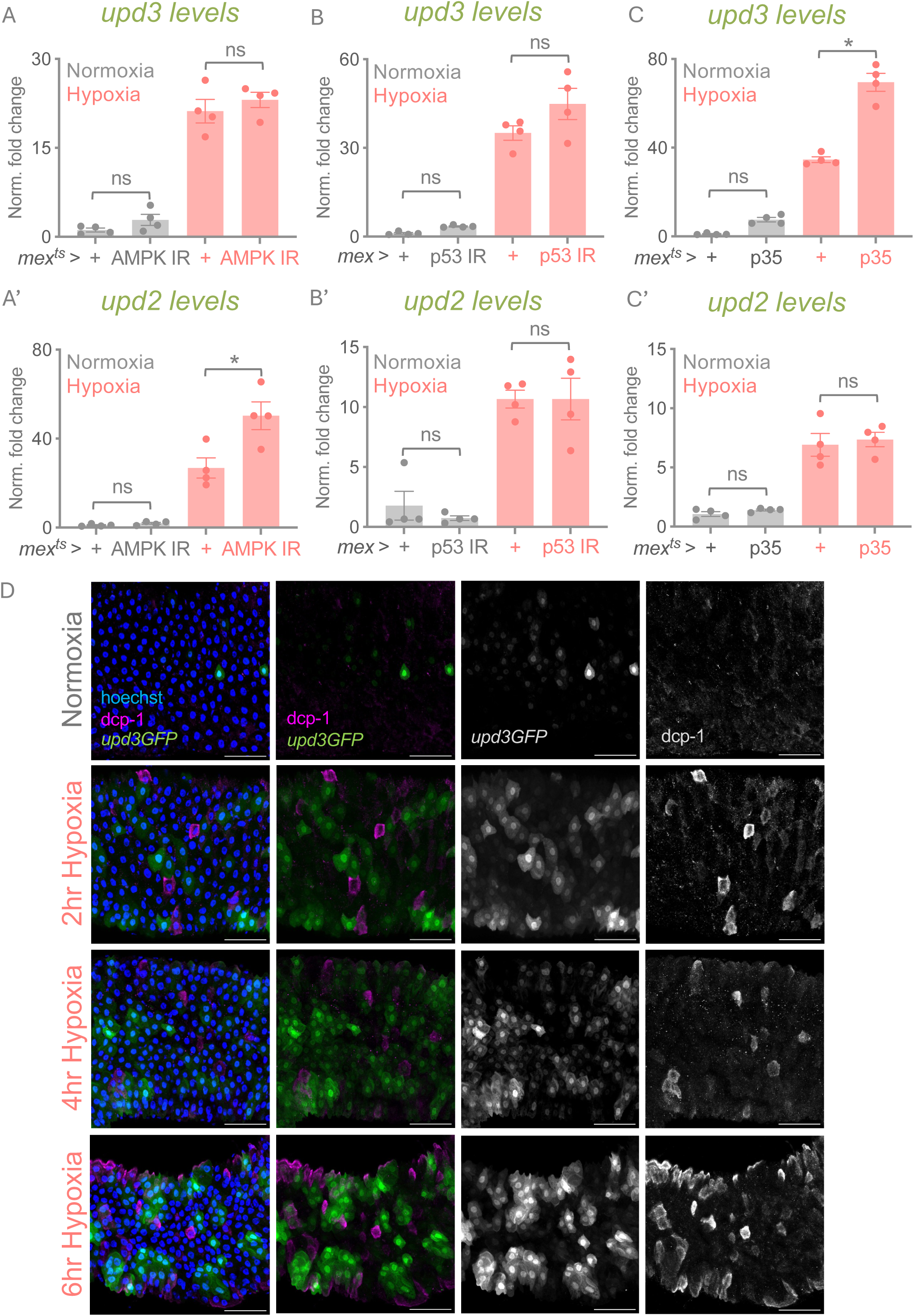
Upd cytokine induction in hypoxia is independent of the AMPK-p53-caspase pathway. (A-C, A’-C’) *upd2* and *upd3* induction in hypoxia is independent of the pathway driving enterocyte apoptosis. Blocking this pathway by enterocyte-specific expression of (A) dsRNA to *AMPK*, (B) dsRNA to *p53*, or (C) p35 resulted in no significant suppression of *upd2* and *upd3* transcript levels induced in hypoxia. Data represents mean ± SEM, N=4 independent groups of samples (5 animals per group) per experimental condition. *p<0.05, ns: not significant (two-way ANOVA with Tukey’s multiple comparisons test). (D) Visualizing *upd3* reporter expression, and caspase-positive cells simultaneously indicates that caspase-inducing cells are largely separable from Upd3 producing cells in early (2 hours), mid (4 hours), or late (6 hours) hypoxia. Scale bar = 50 *μ*m.

We also examined whether hypoxic Upd induction occurs through signaling pathways previously implicated in injury-induced Upd production. The best-characterized of these are ROS-dependent activation of the stress kinases JNK and p38, together with Hippo-Yorkie signaling, each defined in the context of pathogen infection, ingested toxins or DNA-damaging agents (Houtz *et al*., 2017; Jiang *et al*., 2009; Li *et al*., 2014; Mundorf *et al*., 2019; Patel *et al*., 2019; Santabarbara-Ruiz *et al*., 2015; Staley & Irvine, 2010; Zhou *et al*., 2013). We have previously shown that hypoxic Upd induction in the gut is independent of ROS (Ding *et al*., 2026). To test the remaining pathways, we used <u>the enterocyte-specific driver mex-Gal4</u> to express a dominant-negative form of Basket (Bsk), the *Drosophila* JNK, or RNAi against *p38b* or *yorkie*. None of these manipulations suppressed hypoxia-induced *upd2* and *upd3* expression, and several increased it (Fig. S3). Hypoxic Upd induction is therefore independent of the pathways most strongly implicated in Upd production following infection or epithelial damage.

Together these findings indicate that hypoxia induces two parallel responses in largely separate enterocyte populations, one activating an AMPK-p53-*reaper* pathway that primes cells for apoptosis and the other inducing Upd cytokines that prime stem cells for proliferation. Both are initiated during hypoxia, and neither is completed until oxygen is restored, coupling epithelial cell loss and stem cell-mediated repair to the reoxygenation phase.

## Discussion

Fluctuations in oxygen availability are a recurring feature of intestinal physiology and are further amplified in many disease states (Dvornikova *et al*., 2023; Singhal & Shah, 2020; Solanki & Shah, 2024; Villareal & Xue, 2024; Zheng *et al*., 2015). How epithelial tissues adapt to these dynamic oxygen environments while maintaining tissue integrity, however, remains poorly understood. Using the adult *Drosophila* intestine, we identify a two-stage response to hypoxia-reoxygenation in which hypoxia initiates parallel enterocyte responses that are executed only when oxygen is restored. During hypoxia, one enterocyte population activates an AMPK-p53-dependent response that primes cells for apoptosis, while a second, largely non-overlapping population induces Upd cytokine signaling independently of this arm, priming intestinal stem cells for regeneration. Upon reoxygenation, both are completed, and enterocyte loss and stem cell-mediated regeneration proceed together. Our findings therefore suggest that adaptation to fluctuating oxygen availability involves the separation of stress and regenerative signaling across enterocyte populations, and a delay between the initiation of these responses and their execution.

One question raised by these results is why execution of both responses is deferred until reoxygenation. Enterocytes activate caspases and progenitors activate JAK-STAT signaling during hypoxia, yet neither apoptosis nor ISC proliferation occurs until oxygen is restored. One possibility is that low oxygen limits the energetic or biosynthetic capacity needed to complete apoptosis and cell division. Consistent with this, the caspase response requires AMPK, whose activation reports a fall in cellular energy charge. On this view the same deficit that primes a cell for apoptosis is what prevents it from completing it, and both are resolved on reoxygenation. A second is that low oxygen induces a response that actively restrains execution until oxygen returns. Alternatively, reoxygenation itself may supply an instructive signal that triggers completion of apoptosis and induction of ISC proliferation. Intestinal ischemia-reperfusion offers a precedent for this logic, since restoring blood flow after an ischemic episode causes epithelial injury driven in large part by a burst of reactive oxygen species upon the sudden return of oxygen (Archontakis-Barakakis *et al*., 2025; Goncharov & Sharapov, 2023; Wu *et al*., 2018). ROS are established upstream regulators of both apoptosis (Buchon *et al*., 2009b; Jiang *et al*., 2009; Takeishi *et al*, 2013) and ISC proliferation (Hochmuth *et al*, 2011; Perochon *et al*., 2021; Tamamouna *et al*., 2021; Wang *et al*, 2014) in the fly intestine, raising the possibility that a comparable burst contributes here. More generally, a similar temporal organization is observed during nutrient deprivation, where enterocyte Upd production and progenitor JAK-STAT activation occur during starvation, but the proliferative response emerges only after refeeding (Obata *et al*, 2018). The parallels between these systems suggest that the intestine may employ a common adaptive strategy in which adverse environmental conditions initiate a regenerative response while its downstream consequences are completed only once conditions improve.

Our data show that the two enterocyte responses, although induced separately during hypoxia, are coupled in the control of ISC proliferation at reoxygenation. Proliferation is blunted when EC caspases are inhibited and when Upd-STAT signaling is removed. One possibility is that ISC proliferation initiates only while enterocytes are actively being lost, whether because dying cells release a signal or because their removal lifts a constraint, so that the EC replacement is tightly coupled to EC loss (Liang *et al*, 2017; Loudhaief *et al*, 2017; O’Brien & Bilder, 2013). A comparable mechanism operates during homeostatic turnover in this tissue, where dying enterocytes trigger EGF-dependent division of neighboring stem cells (Ngo *et al*, 2020). In this view, the absence of proliferation during hypoxia, despite STAT activation in progenitors, would follow at least in part from the pause in apoptosis of AMPK-p53-primed enterocytes. A direct restraint on ISC division by the hypoxic environment may also contribute, and the two need not be alternatives since either alone would be sufficient to hold division in check until reoxygenation.

A further question raised by our findings is why individual enterocytes mount different responses to the same hypoxic episode. Each response is induced in only a subset of enterocytes, and the two occur largely, though not exclusively, in different cells. One possibility, consistent with the energetic account above, is that response identity reflects the severity of deprivation experienced by individual cells. Because oxygen is delivered through the tracheal system, local availability is likely to vary across the epithelium. Cells farthest from tracheal terminal branches would sustain the greatest energetic deficit, and so would be the most likely to activate AMPK, an energy sensor, and be primed for removal, whereas cells under milder limitation would instead induce Upd. Alternatively, the responses may be determined by intrinsic properties of individual enterocytes. Enterocyte age is one potential determinant (Qin *et al*, 2025), as older enterocytes are known to be more susceptible to damage-induced elimination during pathogenic infection (Akagi *et al*, 2018; Loudhaief *et al*., 2017; Nagy & Buchon, 2025). More broadly, recent single-cell analyses have revealed substantial transcriptional and physiological heterogeneity within the enterocyte compartment (Dutta *et al*, 2015; Hung *et al*, 2020; Hung *et al*, 2021; Li *et al*, 2022; Shin *et al*, 2022). In this view, hypoxia would not generate distinct enterocyte states de novo but would act on pre-existing differences between cells to bias them toward one response or the other.

The requirement for Upd signaling observed here is consistent with its established role in intestinal regeneration following pathogen infection, ingested toxins and DNA-damaging agents, where enterocyte-derived Upd cytokines stimulate the ISC proliferation that restores epithelial integrity (Beebe *et al*., 2010; Jiang *et al*., 2009; Zhou *et al*., 2013). What appears different in the hypoxia response is the relationship between stress sensing and Upd induction. In previously characterized injury models, the pathways that detect epithelial damage also promote *upd* expression, directly coupling stress detection to regenerative signaling (Jiang *et al*., 2009; Mundorf *et al*., 2019; Patel *et al*., 2019; Santabarbara-Ruiz *et al*., 2015; Staley & Irvine, 2010; Zhang & Edgar, 2022). In contrast, our data suggest that hypoxia-induced Upd production is uncoupled from the pathway that senses hypoxic stress. Upd induction occurs independently of the AMPK-p53 pathway that drives apoptotic priming, and hypoxic *upd* expression remains intact following inhibition of JNK, p38 and Hippo/Yki signaling. We previously found that *upd3* induction is also independent of ROS and of Sima, the canonical mediator of oxygen-sensing and an obvious candidate in this context (Ding *et al*., 2026).

What induces Upd under hypoxia therefore remains an open question. One possibility is mechanical signaling, as deformation has been shown to drive *upd* expression in this tissue (Li *et al*, 2018), and changes in enterocyte size and shape represent the earliest response we detect, coinciding with the onset of Upd induction. Why the stress pathway activated by hypoxia in enterocytes does not itself account for Upd induction is likewise unclear and may reflect the nature of the perturbation. An emerging view in tissue biology is that cellular responses are shaped not simply by the presence of stress, but by the specific type of disturbance imposed, whether a loss of structure, function or homeostatic control (Medzhitov, 2021).

Our findings with Sima further emphasize how tissue-level adaptation to hypoxia-reoxygenation involves distinct responses across intestinal cell types. We find that hypoxia induces Sima activity within the intestine, but this activity is most pronounced in progenitor cells rather than enterocytes. Its functional requirement follows the same pattern, with *sima* dispensable for the enterocyte responses described here but required in progenitor cells for the proliferative response that follows reoxygenation. Because *sima* induction occurs during hypoxia, one possibility is that Sima acts at that time, as part of the priming of progenitors. Alternatively, *sima* may be required during reoxygenation to support ISC proliferation. In this context it may function similarly to its pro-proliferative role in cancer cells, where HIF-1α promotes division through the induction of metabolic genes, including *Ldh*, which we also find induced during hypoxia (DeBerardinis *et al*, 2008; Rankin *et al*, 2016; Schito & Semenza, 2016; Xie & Simon, 2017). Consistent with a requirement at that stage, ROS and HIF-1α reinforce one another to promote stem cell proliferation in the distal mouse colon (Chen *et al*, 2025), indicating that redox state and HIF-1α activity can act together to control proliferation in an intestinal epithelium. A further possibility is that Sima acts together with STAT, since both are required in progenitors for the same divisions. Transcriptional cooperation between HIF-1α and STAT3 has been described in zebrafish tissues and regenerating mouse skin epithelia (Dinarello *et al*, 2023; Liu *et al*, 2023).

Collectively, our findings indicate that adaptation to fluctuating oxygen availability emerges from the coordinated behavior of multiple cell populations rather than a response shared by all cells, with hypoxia initiating these responses and reoxygenation permitting their completion. The importance of this organization is underscored by the reduced survival observed when the enterocyte response is disrupted during intermittent hypoxia. Neither phase alone is sufficient to explain the resulting tissue behavior, suggesting that the tissue responds to oxygen fluctuation rather than to hypoxia or reoxygenation in isolation. Because such cycles occur during normal intestinal physiology and are amplified in ischemia-reperfusion injury, inflammatory bowel disease and sleep-disordered breathing, the organizational principles defined here may extend beyond the Drosophila intestine.

## Methods

### Fly Stocks and Husbandry

*Drosophila* stocks were maintained at 25°C or 18°C. All stocks were maintained on media containing 100g *Drosophila* Type II agar, 1200g cornmeal, 490g Torula yeast, 450g sugar, 1240g D-glucose, 160mL acid mixture (propionic and phosphoric acid) in 20L water. Genetic crosses were established by mating virgin females with males. All crosses and progeny were maintained at 25°C, or 18°C (for temperature-sensitive GAL4 crosses).

The following stocks were used in this study: *Myo1A-Gal4^ts^* (Jiang *et al*., 2009), *mex-Gal4* [Bloomington Drosophila Stock Center (BDSC) 91368], *mex-Gal4^ts^*, *esg-Gal4^ts^* (gifted from Bruce Edgar, Huntsman Cancer Institute, University of Utah, Salt Lake City, UT, USA), *UAS-p35* (BDSC 5072), *UAS-AMPK^DN^* (BDSC 32112), *UAS-AMPK^CA^* (BDSC 32110), *UAS-AMPKα RNAi* [Vienna *Drosophila* Resource Center (VDRC) 1827/GD], *UAS-p53 RNAi* (VDRC 38235), *UAS-STAT92E RNAi* (VDRC 43866/GD), *UAS-sima RNAi* (VDRC 106187/KK), *10XSTAT-GFP, upd3-Gal4, UAS-GFP* (gifted from Erica Bach, NYU Grossman School of Medicine, New York, NY, USA), *upd2,3Δ* (BDSC 55729), *FRT82B* (BDSC 86313), *MARCM82B* (gifted from Bruce Edgar), *UAS-yki RNAi* (BDSC 34067), *UAS-p38b RNAi* (BDSC 29405), *UAS-bsk^DN^* (BDSC 6409). We used FlyBase (FB 2026_01) to find information on all genes and fly stocks used in this study (Jenkins *et al*, 2022; Ozturk-Colak *et al*, 2024). See Table S1 for complete list of lines used.

### Hypoxia Exposure

Adult Drosophila vials were placed in an airtight chamber, with a constant flow of 1% oxygen (1% oxygen/99% nitrogen) at 23°C. Flow rate through the chamber was regulated using an Aalborg model P gas flow meter. Normoxic control animals were handled identically and held at the same temperatures for time periods matching the hypoxia and reoxygenation exposures.

### Adult-specific Gene Expression

Mated female flies were used for all experiments. Adult-specific transgene expression was achieved using *tubGal80^ts^*, by raising flies at 18°C throughout development until 1 week after eclosion. Flies were then shifted to 29°C to induce GAL4-mediated gene expression under the control of cell-type specific drivers. Transgene induction was carried out for 2 days prior to hypoxia exposure. For reoxygenation timepoints, flies were moved back to 29°C following hypoxia exposure to maintain transgene expression.

### Clonal analysis

MARCM clones were induced in 1-week-old mated female flies by heat shock at 37°C for 20 minutes in a water bath. Flies were then returned to 23°C. For the experiments in Fig. 2C–D, clones were allowed to grow for one week before hypoxia exposure, reaching 6-10 cells per clone. For the experiments in Fig. 3A–B, flies were exposed to hypoxia immediately after heat shock, so that clones began as single cells. Clone size was scored as the number of GFP-positive cells per clone.

### Tissue Dissection and Microscopy

Adult female flies were rapidly surface sterilized in 75% ethanol and transferred to ice cold 1xPBS in a glass dissection dish. Whole guts were dissected using fine forceps under a light microscope and immediately moved to a 9-well dissection plate containing 4% PFA (Electron Microscopy Sciences) for 1 hour at RT. Midguts were then quick washed with 1xPBS three times and incubated with 1xPAT blocking solution (PBS-T containing 10% BSA) for 2 hours at RT. Primary antibodies were prepared in blocking solution and tissues were then incubated overnight at 4°C. The next day, tissues were washed three times for 15 min with 1xPAT and incubated with secondary antibody for 2 hours at RT. Finally, tissues were washed three times for 15 mins with 1xPAT, with the second wash containing Hoechst 33342 nuclear stain (1:1000 dilution). Midguts were then mounted onto glass slides with spacers in Vectashield mounting medium (Vector Laboratories Inc.). Slides were imaged using a Zeiss LSM 880 laser confocal microscope with 10x and 20x objectives and ZEN software.

Primary antibodies were used with the following dilutions: Rabbit anti-phospho-Histone H3 (pSer^10^) [1:500, Millipore Sigma #06-570], Rabbit anti-cleaved-*Drosophila*-Dcp1 (Asp215) [1:500, Cell Signaling Technology, #9578, RRID: AB_2721060], Mouse anti-Discs-large (Dlg) [1:30, Developmental Studies Hybridoma Bank, #4F3, RRID: AB_528203], Phalloidin labelling probes AF488, AF568, [1:1000, Invitrogen (ThermoFisher), #A12379, #A12380]. Secondary antibodies were used with the following dilutions: Goat-anti-rabbit Alexa Fluor 568 [1:2000, Invitrogen (ThermoFisher), #A-11011], Goat-anti-mouse Alexa Fluor 568 [1:2000, Invitrogen (ThermoFisher), #A21124], Goat-anti-mouse Alexa Fluor 488 [1:2000, Invitrogen (ThermoFisher), #A11001], Streptavidin-Alexa Fluor 568 conjugate [1:2000, Invitrogen (ThermoFisher), #S11226].

All images were taken in the anterior midgut in the R2 region (Buchon & Osman, 2015; Buchon *et al*, 2013; Marianes & Spradling, 2013; O’Brien, 2013). Images were processed using Fiji (Schindelin *et al*, 2012).

### Gut length measurements

Whole guts were dissected as described above and imaged by brightfield microscopy. Midgut length was measured in Fiji as the distance between the anterior and posterior landmarks indicated by arrowheads in Fig. 1A. For each experiment, lengths were normalized to the mean of the corresponding normoxic control group.

### TUNEL assay

Adult tissues were dissected and fixed as described above. Tissues were then permeabilized in PBS-T (1xPBS containing 0.3% Triton-X) for 15 mins. Fixed, permeabilized intestines were then treated with DeadEnd TUNEL Labelling system (Promega G7360) following manufacturer’s instructions. TUNEL detection was carried out using Streptavidin-AF568 secondary antibody.

### qRT-PCR

Female gut tissues were dissected in ice cold 1xPBS in groups of 5 and immediately transferred to 1.5mL microcentrifuge tubes containing 500 *μL* TRIzol reagent (Invitrogen #15596-018) before snap freezing on dry ice. Total RNA was extracted from adult tissues following manufacturer’s instructions (Invitrogen #15596-018). RNA samples were then DNAse treated following manufacturer’s instructions (Ambion, #2238 G). Reverse transcription was achieved using SuperScript III (Invitrogen, #18080044). Generated cDNA was used as a template for qRT-PCR reactions (QuantStudio 6 RT-PCR system, Applied Biosystems) using the primer pairs defined in Table S2, and SYBR Green reagents (Applied Biosystems, Foster City, CA, USA). Data analysis for qRT-PCR was completed using the comparative CT method (2^-ΔΔCT^). PCR data were normalized to reference genes (*5S rRNA*, *rps9* mRNA or *eif2alpha* mRNA) whose expression levels were unaffected by experimental conditions.

### Statistics

For all experiments, error bars represent mean ± SEM. P-values are from unpaired two-tailed t-tests, or one-way or two-way ANOVA with Tukey’s multiple comparisons test, or Mantel-Cox log rank test for survival, calculated in GraphPad Prism version 11.1.0 (GraphPad Software, Boston, Massachusetts USA). P<0.05 was considered statistically significant.

## Acknowledgements

We thank Bruce Edgar for the *mex*-Gal4^ts^, *esg*-Gal4^ts^ and MARCM82B lines, Erica Bach for the 10XSTAT-GFP, *upd3*-Gal4 and UAS-GFP lines, and Irene Miguel-Aliaga for the *ldh::gfp* line. We thank the Bloomington Drosophila Stock Center (NIH P40OD018537) and the Vienna Drosophila Resource Center for fly stocks, and the Developmental Studies Hybridoma Bank, created by the NICHD of the NIH and maintained at the University of Iowa, for antibodies. This work was supported by the Canadian Institutes of Health Research (Project Grants PJT-173517 and PJT-195892) and the Natural Sciences and Engineering Research Council of Canada (Discovery Grant RGPIN-2023-04524) to S.S.G.

## Author Contributions

P.B. and S.S.G. conceived and designed the study. P.B. performed experiments and analyzed the data. K.D. designed and performed experiments and analyzed data. P.B. and S.S.G. wrote the original draft, and S.S.G. revised the manuscript. S.S.G. provided supervision and funding.

---

**Figure S1:**
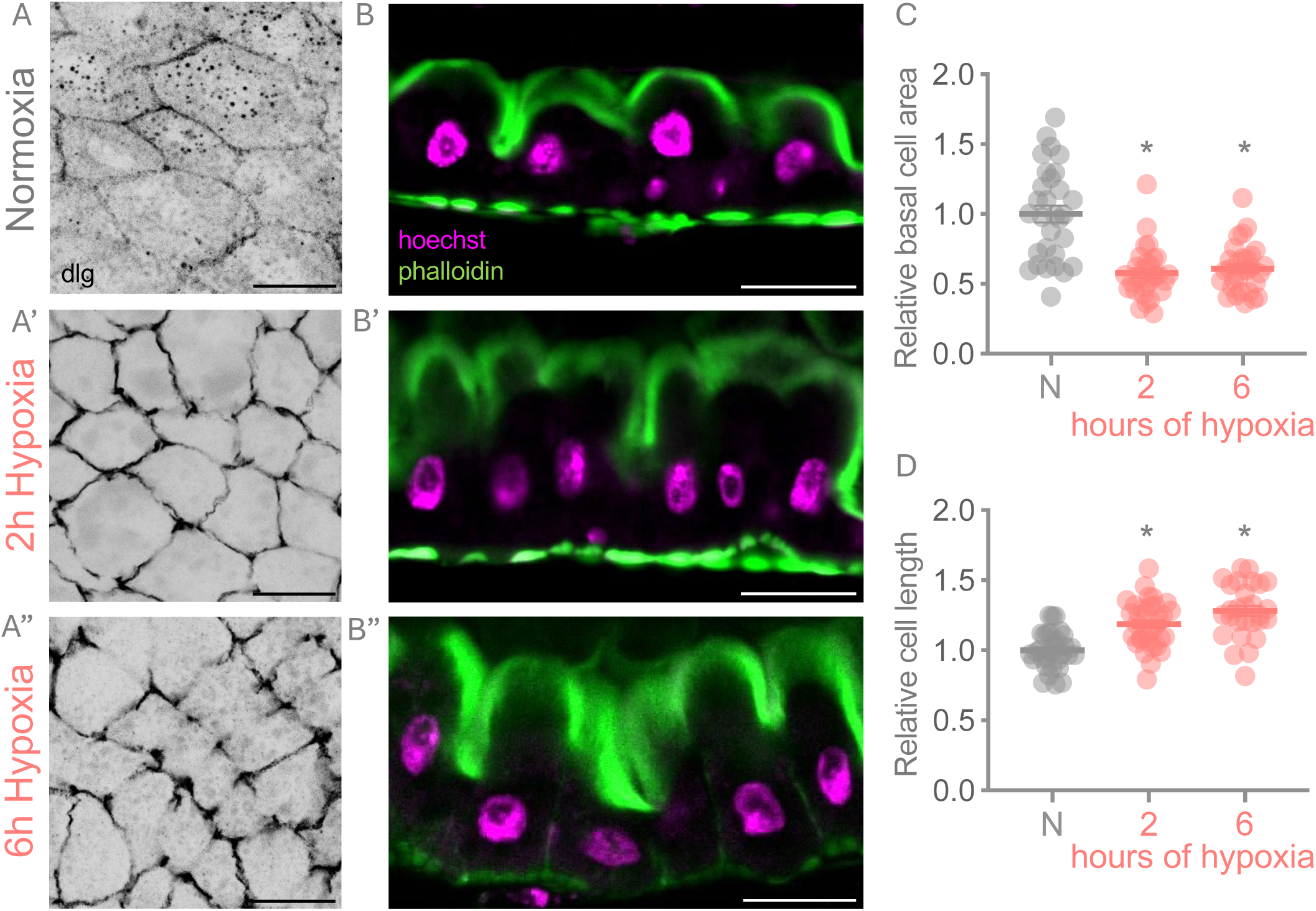
Acute hypoxia alters epithelial enterocyte morphology. (A-A”) Hypoxic guts show reduced basal cell area as seen by confocal images of guts stained for Discs large (dlg) to visualize basal cell boundaries per experimental condition in the R2 region. Scale bar = 20*μ*m. (B-B”) Confocal images of guts stained with Hoechst nuclear stain and Phalloidin to visualize cell apical-basal length in the R2 region show increased cell length in hypoxic intestines. Scale bar = 20*μ*m. Quantification of relative cell area (C) and relative cell length (D) in normoxia, compared to 2, or 6 hours of hypoxia (1% Oxygen). n = 28-30 cells per experimental group (5-10 independent tissues). *p<0.05 (unpaired t-test).

**Figure S2:**
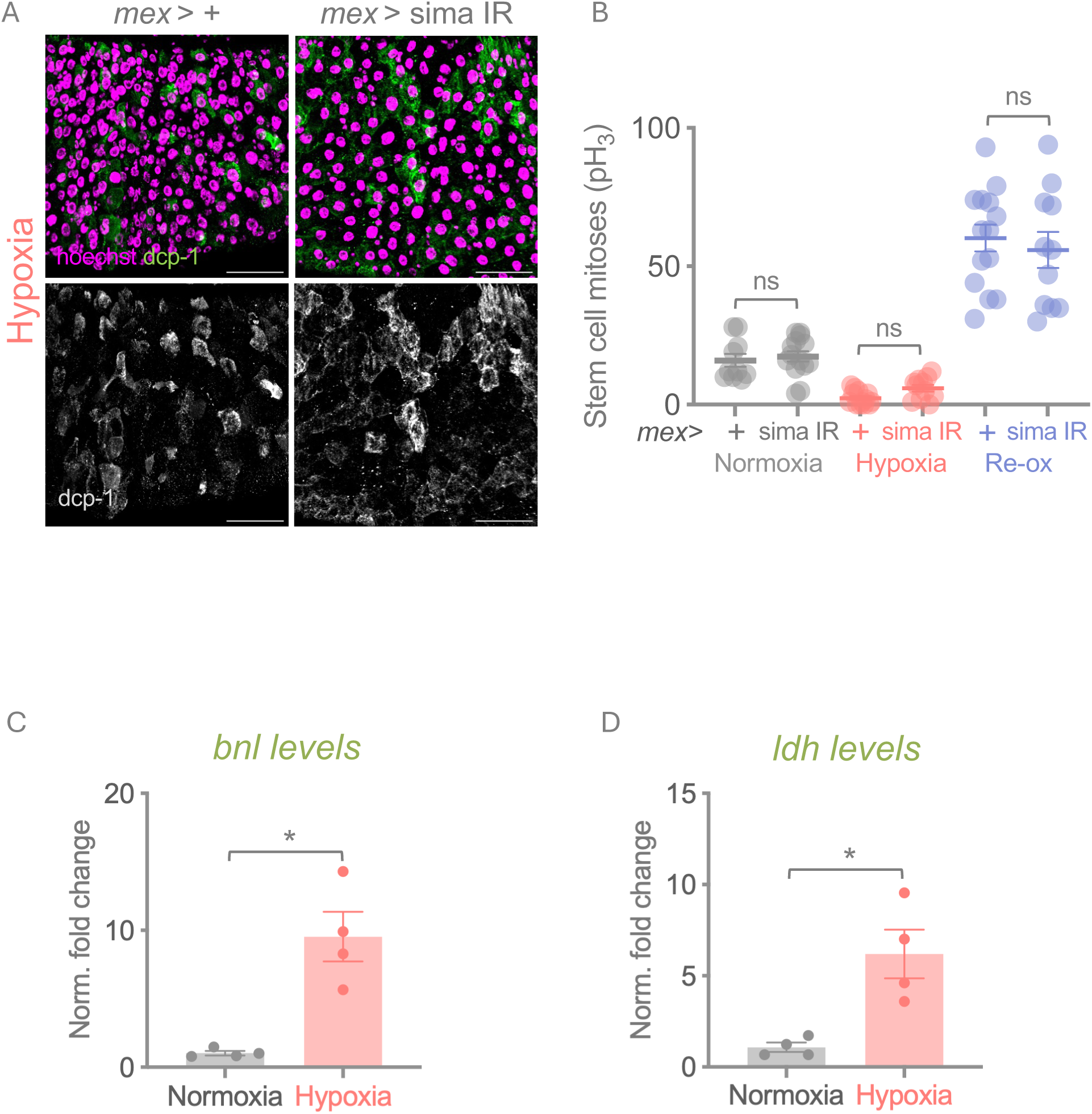
Enterocyte HIF-1α is not required for caspase induction in hypoxia. (A) Guts were stained for Dcp-1 to measure caspase activity in intestines exposed to hypoxia. Control intestines induce caspase activity in large polyploid cells. Expression of dsRNA to HIF-1α in intestinal enterocytes, resulted in no change in caspase induction during the hypoxic period. Scale bar = 50 *μ*m.(B) Expression of dsRNA to HIF-1α in enterocytes has no impact on stem cell divisions under normoxia, hypoxia, or reoxygenation. Each data point depicts total proliferating stem cells in individual tissues and data represents mean ± SEM, n = 10-15 guts per experimental condition. ns: not significant (two-way ANOVA with Tukey’s multiple comparisons test). (C, D) Sima target genes *branchless (bnl)* and *lactate dehydrogenase (ldh)* transcripts are upregulated in hypoxic intestines. Data represents mean ± SEM, N=4 independent groups of samples (5 animals per group) per experimental condition. *p<0.05 (unpaired t-test). Data points represent independent samples.

**Figure S3:**
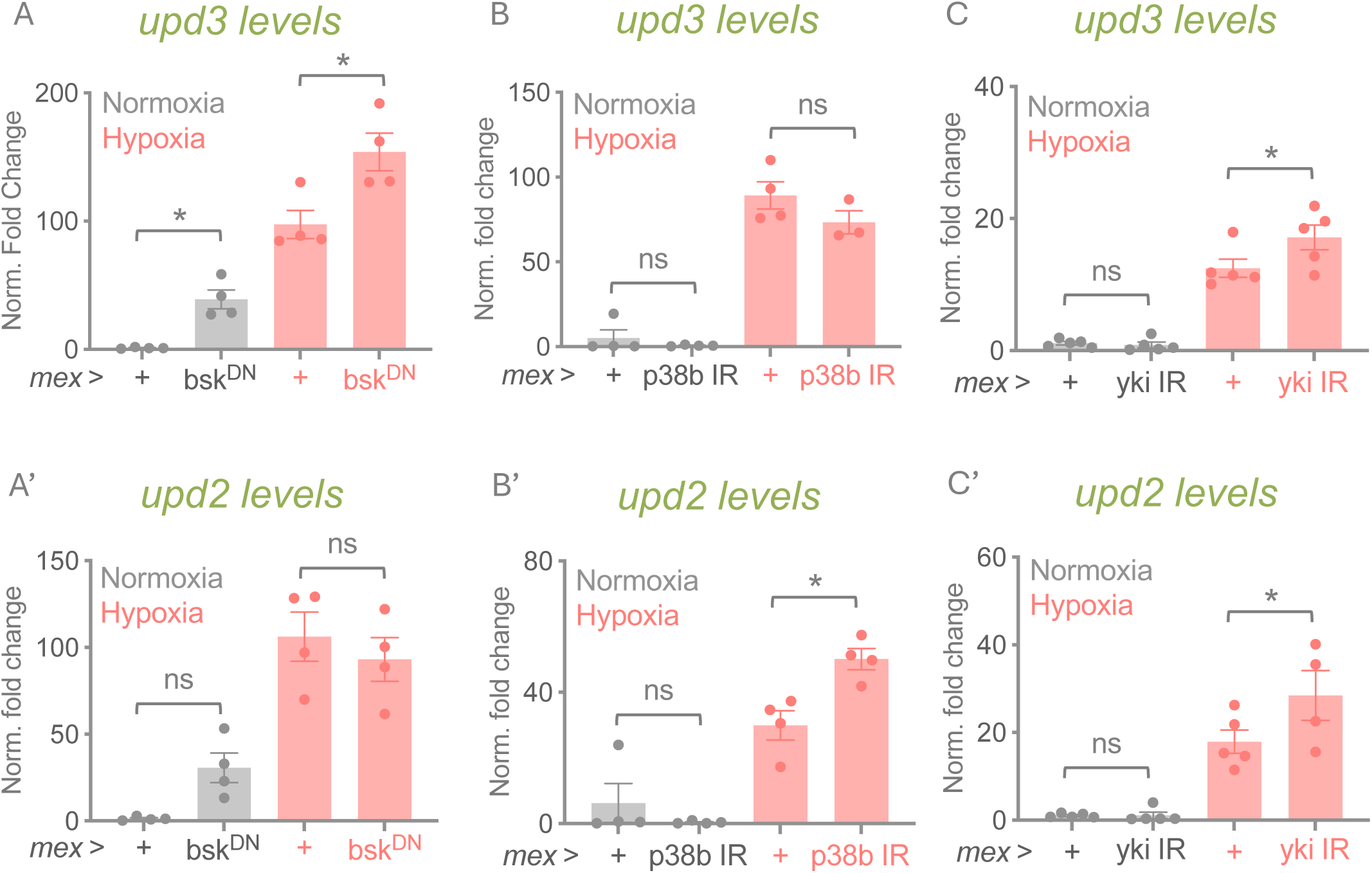
*Unpaired* cytokine induction in hypoxia is not dependent on JNK, p38, and Hpo/Yki signaling. (A-C) *upd3* transcript levels were measured in isolated intestines from animals exposed to normoxia or hypoxia. *upd3* induction in hypoxia showed no significant suppression compared to hypoxic controls under enterocyte-specific expression of (A) *bsk^DN^*, (B) dsRNA to *p38b*, or (C) dsRNA to *yki*. Data represents mean ± SEM, N=4 independent groups of samples (5 animals per group) per experimental condition. *p<0.05, ns: not significant (two-way ANOVA with Tukey’s multiple comparisons test). Data points represent independent samples. (A’-C’) *upd2* transcript levels were measured in isolated intestines from animals exposed to normoxia or hypoxia. *upd2* induction in hypoxia showed no significant suppression compared to hypoxic controls under enterocyte-specific expression of (A’) *bsk^DN^*, (B’) dsRNA to *p38b*, or (C’) dsRNA to *yki*. Data represents mean ± SEM, N=4 independent groups of samples (5 animals per group) per experimental condition. *p<0.05, ns: not significant (two-way ANOVA with Tukey’s multiple comparisons test). Data points represent independent samples.

## References

Akagi K, Wilson K, Katewa S, Ortega M, Simons J, Hilsabeck T, Kapuria S, Sharma A, Jasper H, Kapahi P (2018) Dietary restriction improves intestinal cellular fitness to enhance gut barrier function and lifespan in D-melanogaster. Plos Genetics 14

Archontakis-Barakakis P, Mavridis T, Chalkias A (2025) Oxygen-Mediated Molecular Mechanisms Involved in Intestinal Ischemia and Reperfusion Injury. International Journal of Molecular Sciences 26

Bach EA, Ekas LA, Ayala-Camargo A, Flaherty MS, Lee H, Perrimon N, Baeg GH (2007) GFP reporters detect the activation of the Drosophila JAK/STAT pathway in vivo. Gene Expr Patterns 7: 323–331

Beebe K, Lee W, Micchelli C (2010) JAK/STAT signaling coordinates stem cell proliferation and multilineage differentiation in the Drosophila intestinal stem cell lineage. Developmental Biology 338: 28–37

Beumer J, Clevers H (2016) Regulation and plasticity of intestinal stem cells during homeostasis and regeneration. Development 143: 3639–3649

Beumer J, Clevers H (2021) Cell fate specification and differentiation in the adult mammalian intestine. Nat Rev Mol Cell Biol 22: 39–53

Biteau B, Hochmuth C, Jasper H (2008) JNK Activity in Somatic Stem Cells Causes Loss of Tissue Homeostasis in the Aging Drosophila Gut. Cell Stem Cell 3: 442–455

Biteau B, Hochmuth CE, Jasper H (2011) Maintaining tissue homeostasis: dynamic control of somatic stem cell activity. Cell Stem Cell 9: 402–411

Blackie L, Gaspar P, Mosleh S, Lushchak O, Kong L, Jin Y, Zielinska AP, Cao B, Mineo A, Silva B et al (2024) The sex of organ geometry. Nature 630: 392–400

Boumard B, Bardin A (2021) An amuse-bouche of stem cell regulation: Underlying principles and mechanisms from adult Drosophila intestinal stem cells. Current Opinion in Cell Biology 73: 58–68

Brodsky M, Nordstrom W, Tsang G, Kwan E, Rubin G, Abrams J (2000) Drosophila p53 binds a damage response element at the reaper locus. Cell 101: 103–113

Brown E, Taylor CT (2018) Hypoxia-sensitive pathways in intestinal inflammation. J Physiol 596: 2985–2989

Buchon N, Broderick NA, Chakrabarti S, Lemaitre B (2009a) Invasive and indigenous microbiota impact intestinal stem cell activity through multiple pathways in Drosophila. Genes Dev 23: 2333–2344

Buchon N, Broderick NA, Poidevin M, Pradervand S, Lemaitre B (2009b) Drosophila intestinal response to bacterial infection: activation of host defense and stem cell proliferation. Cell Host Microbe 5: 200–211

Buchon N, Osman D (2015) All for one and one for all: Regionalization of the Drosophila intestine. Insect Biochemistry and Molecular Biology 67: 2–8

Buchon N, Osman D, David F, Fang H, Boquete J, Deplancke B, Lemaitre B (2013) Morphological and Molecular Characterization of Adult Midgut Compartmentalization in Drosophila. Cell Reports 3: 1725–1738

Centanin L, Dekanty A, Romero N, Irisarri M, Gorr TA, Wappner P (2008) Cell autonomy of HIF effects in Drosophila: tracheal cells sense hypoxia and induce terminal branch sprouting. Dev Cell 14: 547–558

Chen X, Raghunathan K, Bao B, Ngwa E, Kwong A, Wu Z, Babcock S, Baek C, Ye G, Muppirala A et al (2025) A niche-dependent redox rheostat regulates epithelial stem cell fate. Nat Commun 16: 11713

Colgan S, Campbell E, Kominsky D (2016) Hypoxia and Mucosal Inflammation. In: Annual Review of Pathology: Mechanisms of Disease, Vol 11, Abbas A.K., Aster J.C., Galli S.J. (eds.) pp. 77–100.

Colgan SP, Taylor CT (2010) Hypoxia: an alarm signal during intestinal inflammation. Nat Rev Gastroenterol Hepatol 7: 281–287

Colombani J, Andersen DS (2020) The Drosophila gut: A gatekeeper and coordinator of organism fitness and physiology. Wiley interdisciplinary reviews Developmental biology 9: e378

DeBerardinis RJ, Lum JJ, Hatzivassiliou G, Thompson CB (2008) The biology of cancer: metabolic reprogramming fuels cell growth and proliferation. Cell metabolism 7: 11–20

Dengler F (2020) Activation of AMPK under Hypoxia: Many Roads Leading to Rome. International Journal of Molecular Sciences 21

Dichtel-Danjoy M, Ma D, Dourlen P, Chatelain G, Napoletano F, Robin M, Corbet M, Levet C, Hafsi H, Hainaut P et al (2013) Drosophila p53 isoforms differentially regulate apoptosis and apoptosis-induced proliferation. Cell Death and Differentiation 20: 108–116

Dinarello A, Betto RM, Diamante L, Tesoriere A, Ghirardo R, Cioccarelli C, Meneghetti G, Peron M, Laquatra C, Tiso N et al (2023) STAT3 and HIF1alpha cooperatively mediate the transcriptional and physiological responses to hypoxia. Cell Death Discov 9: 226

Ding K, Bodkhe P, Lee B, Ingelson-Filpula W, Polan D, Wycislik A, Cheung T, Wu S, Rahim R, Grewal S (2026) Gut-derived Unpaired-3 cytokine signaling promotes systemic hypoxia tolerance in Drosophila. Disease Models & Mechanisms 19

Dutta D, Dobson A, Houtz P, Glasser C, Revah J, Korzelius J, Patel P, Edgar B, Buchon N (2015) Regional Cell-Specific Transcriptome Mapping Reveals Regulatory Complexity in the Adult Drosophila Midgut. Cell Reports 12: 346–358

Dvornikova K, Platonova O, Bystrova E (2023) Hypoxia and Intestinal Inflammation: Common Molecular Mechanisms and Signaling Pathways. International Journal of Molecular Sciences 24

Ghoshal S, Chard E, Shore S, Hartman J, Engevik M (2026) OXYGEN DEPLETION BY GUT MICROBES AS A REGULATOR OF INTESTINAL INFLAMMATION. Gastroenterology 170

Goncharov R, Sharapov M (2023) Ischemia-Reperfusion Injury: Molecular Mechanisms of Pathogenesis and Methods of Their Correction. Molecular Biology 57: 1143–1164

Guo Z, Ohlstein B (2015) Bidirectional Notch signaling regulates Drosophila intestinal stem cell multipotency. Science 350

Hageman JH, Heinz MC, Kretzschmar K, van der Vaart J, Clevers H, Snippert HJG (2020) Intestinal Regeneration: Regulation by the Microenvironment. Dev Cell 54: 435–446

Hardie D, Ross F, Hawley S (2012) AMPK: a nutrient and energy sensor that maintains energy homeostasis. Nature Reviews Molecular Cell Biology 13: 251–262

Hay B, Wolff T, Rubin G (1994) EXPRESSION OF BACULOVIRUS P35 PREVENTS CELL-DEATH IN DROSOPHILA. Development 120: 2121–2129

Hochmuth C, Biteau B, Bohmann D, Jasper H (2011) Redox Regulation by Keap1 and Nrf2 Controls Intestinal Stem Cell Proliferation in Drosophila. Cell Stem Cell 8: 188–199

Hou SX (2010) Intestinal stem cell asymmetric division in the Drosophila posterior midgut. J Cell Physiol 224: 581–584

Houtz P, Bonfini A, Liu X, Revah J, Guillou A, Poidevin M, Hens K, Huang HY, Deplancke B, Tsai YC et al (2017) Hippo, TGF-beta, and Src-MAPK pathways regulate transcription of the upd3 cytokine in Drosophila enterocytes upon bacterial infection. PLoS Genet 13: e1007091

Hung R, Hu Y, Kirchner R, Liu Y, Xu C, Comjean A, Tattikota S, Li F, Song W, Sui S et al (2020) A cell atlas of the adult Drosophila midgut. Proceedings of the National Academy of Sciences of the United States of America 117: 1514–1523

Hung R, Li J, Liu Y, Perrimon N (2021) Defining cell types and lineage in the Drosophila midgut using single cell transcriptomics. Current Opinion in Insect Science 47: 12–17

Ingaramo MC, Sanchez JA, Perrimon N, Dekanty A (2020) Fat Body p53 Regulates Systemic Insulin Signaling and Autophagy under Nutrient Stress via Drosophila Upd2 Repression. Cell Rep 33: 108321

Jenkins VK, Larkin A, Thurmond J, FlyBase C (2022) Using FlyBase: A Database of Drosophila Genes and Genetics. *Methods in molecular biology (Clifton*, NJ) 2540: 1–34

Jiang H, Edgar BA (2011) Intestinal stem cells in the adult Drosophila midgut. Exp Cell Res 317: 2780–2788

Jiang H, Edgar BA (2012) Intestinal stem cell function in Drosophila and mice. Curr Opin Genet Dev 22: 354–360

Jiang H, Patel PH, Kohlmaier A, Grenley MO, McEwen DG, Edgar BA (2009) Cytokine/Jak/Stat signaling mediates regeneration and homeostasis in the Drosophila midgut. Cell 137: 1343–1355

Jiang H, Tian A, Jiang J (2016) Intestinal stem cell response to injury: lessons from Drosophila. Cellular and molecular life sciences : CMLS 73: 3337–3349

Johnson E, Kazgan N, Bretz C, Forsberg L, Hector C, Worthen R, Onyenwoke R, Brenman J (2010) Altered Metabolism and Persistent Starvation Behaviors Caused by Reduced AMPK Function in Drosophila. Plos One 5

Kelly C, Zheng L, Campbell E, Saeedi B, Scholz C, Bayless A, Wilson K, Glover L, Kominsky D, Magnuson A et al (2015) Crosstalk between Microbiota-Derived Short-Chain Fatty Acids and Intestinal Epithelial HIF Augments Tissue Barrier Function. Cell Host & Microbe 17: 662–671

Lavista-Llanos S, Centanin L, Irisarri M, Russo DM, Gleadle JM, Bocca SN, Muzzopappa M, Ratcliffe PJ, Wappner P (2002) Control of the hypoxic response in Drosophila melanogaster by the basic helix-loop-helix PAS protein similar. Mol Cell Biol 22: 6842–6853

Li H, Janssens J, De Waegeneer M, Kolluru S, Davie K, Gardeux V, Saelens W, David F, Brbic M, Leskovec J et al (2022) Fly Cell Atlas: A single-nucleus transcriptomic atlas of the adult fruit fly. Science 375: 991-+

Li H, Jasper H (2016) Gastrointestinal stem cells in health and disease: from flies to humans. Dis Model Mech 9: 487–499

Li Q, Li S, Mana-Capelli S, Flach R, Danai L, Amcheslavsky A, Nie Y, Kaneko S, Yao X, Chen X et al (2014) The Conserved Misshapen-Warts-Yorkie Pathway Acts in Enteroblasts to Regulate Intestinal Stem Cells in Drosophila. Developmental Cell 31: 291–304

Li Q, Nirala N, Nie Y, Chen H, Ostroff G, Mao J, Wang Q, Xu L, Ip Y (2018) Ingestion of Food Particles Regulates the Mechanosensing Misshapen-Yorkie Pathway in Drosophila Intestinal Growth. Developmental Cell 45: 433-+

Li Z, Zhang Y, Han L, Shi L, Lin X (2013) Trachea-derived dpp controls adult midgut homeostasis in Drosophila. Dev Cell 24: 133–143

Liang J, Balachandra S, Ngo S, O’Brien L (2017) Feedback regulation of steady-state epithelial turnover and organ size. Nature 548: 588-+

Litvak Y, Byndloss M, Bäumler A (2018) Colonocyte metabolism shapes the gut microbiota. Science 362: 1017-+

Liu L, Cash TP, Jones RG, Keith B, Thompson CB, Simon MC (2006) Hypoxia-induced energy stress regulates mRNA translation and cell growth. Mol Cell 21: 521–531

Liu S, Hur YH, Cai X, Cong Q, Yang Y, Xu C, Bilate AM, Gonzales KAU, Parigi SM, Cowley CJ et al (2023) A tissue injury sensing and repair pathway distinct from host pathogen defense. Cell 186: 2127–2143 e2122

Loudhaief R, Brun-Barale A, Benguettat O, Nawrot-Esposito M, Pauron D, Amichot M, Gallet A (2017) Apoptosis restores cellular density by eliminating a physiologically or genetically induced excess of enterocytes in the Drosophila midgut. Development 144: 808–819

Majmundar AJ, Wong WJ, Simon MC (2010) Hypoxia-inducible factors and the response to hypoxic stress. Mol Cell 40: 294–309

Marianes A, Spradling A (2013) Physiological and stem cell compartmentalization within the Drosophila midgut. Elife 2

May AM, Mehra R (2014) Obstructive sleep apnea: role of intermittent hypoxia and inflammation. Semin Respir Crit Care Med 35: 531–544

McCall K, Steller H (1998) Requirement for DCP-1 caspase during Drosophila oogenesis. Science 279: 230–234

Medzhitov R (2021) The spectrum of inflammatory responses. Science 374: 1070–1075

Micchelli CA, Perrimon N (2006) Evidence that stem cells reside in the adult Drosophila midgut epithelium. Nature 439: 475–479

Mollereau B, Ma D (2014) The p53 control of apoptosis and proliferation: lessons from Drosophila. Apoptosis 19: 1421–1429

Mundorf J, Donohoe C, McClure C, Southall T, Uhlirova M (2019) Ets21c Governs Tissue Renewal, Stress Tolerance, and Aging in the Drosophila Intestine. Cell Reports 27: 3019-+

Nagy P, Buchon N (2025) Infectious bacteria, but not the microbiota, induce a NOX-ATM-cytokine pathway that controls epithelial turnover. Cell Rep 44: 116293

Ngo S, Liang J, Su YH, O’Brien LE (2020) Disruption of EGF Feedback by Intestinal Tumors and Neighboring Cells in Drosophila. Curr Biol 30: 1537–1546 e1533

O’Brien L (2013) Regional Specificity in the Drosophila Midgut: Setting Boundaries with Stem Cells. Cell Stem Cell 13: 375–376

O’Brien L, Bilder D (2013) Beyond the Niche: Tissue-Level Coordination of Stem Cell Dynamics. In: Annual Review of Cell and Developmental Biology, *Vol* 29, Schekman R. (ed.) pp. 107–136.

Obata F, Tsuda-Sakurai K, Yamazaki T, Nishio R, Nishimura K, Kimura M, Funakoshi M, Miura M (2018) Nutritional Control of Stem Cell Division through S-Adenosylmethionine in Drosophila Intestine. Dev Cell 44: 741–751 e743

Osman D, Buchon N, Chakrabarti S, Huang YT, Su WC, Poidevin M, Tsai YC, Lemaitre B (2012) Autocrine and paracrine unpaired signaling regulate intestinal stem cell maintenance and division. J Cell Sci 125: 5944–5949

Ozturk-Colak A, Marygold SJ, Antonazzo G, Attrill H, Goutte-Gattat D, Jenkins VK, Matthews BB, Millburn G, Dos Santos G, Tabone CJ et al (2024) FlyBase: updates to the Drosophila genes and genomes database. Genetics 227

Pan D, Hardie D (2002) A homologue of AMP-activated protein kinase in Drosophila melanogaster is sensitive to AMP and is activated by ATP depletion. Biochemical Journal 367: 179–186

Patel P, Dutta D, Edgar B (2015) Niche appropriation by Drosophila intestinal stem cell tumours. Nature Cell Biology 17: 1182-+

Patel P, Penalva C, Kardorff M, Roca M, Pavlovic B, Thiel A, Teleman A, Edgar B (2019) Damage sensing by a Nox-Ask1-MKK3-p38 signaling pathway mediates regeneration in the adult Drosophila midgut. Nature Communications 10

Perochon J, Yu Y, Aughey GN, Medina AB, Southall TD, Cordero JB (2021) Dynamic adult tracheal plasticity drives stem cell adaptation to changes in intestinal homeostasis in Drosophila. Nat Cell Biol 23: 485–496

Qin P, Wang Q, Wu Y, You Q, Li M, Guo Z (2025) Age mosaic of gut epithelial cells prevents aging. Nature Communications 16

Rankin EB, Nam JM, Giaccia AJ (2016) Hypoxia: Signaling the Metastatic Cascade. Trends Cancer 2: 295–304

Ren F, Wang B, Yue T, Yun E, Ip Y, Jiang J (2010) Hippo signaling regulates Drosophila intestine stem cell proliferation through multiple pathways. Proceedings of the National Academy of Sciences of the United States of America 107: 21064–21069

Santabarbara-Ruiz P, Lopez-Santillan M, Martinez-Rodriguez I, Binagui-Casas A, Perez L, Milan M, Corominas M, Serras F (2015) ROS-Induced JNK and p38 Signaling Is Required for Unpaired Cytokine Activation during Drosophila Regeneration. PLoS Genet 11: e1005595

Schindelin J, Arganda-Carreras I, Frise E, Kaynig V, Longair M, Pietzsch T, Preibisch S, Rueden C, Saalfeld S, Schmid B et al (2012) Fiji: an open-source platform for biological-image analysis. Nature Methods 9: 676–682

Schito L, Semenza GL (2016) Hypoxia-Inducible Factors: Master Regulators of Cancer Progression. Trends Cancer 2: 758–770

Shaw R, Kohlmaier A, Polesello C, Veelken C, Edgar B, Tapon N (2010) The Hippo pathway regulates intestinal stem cell proliferation during Drosophila adult midgut regeneration. Development 137: 4147–4158

Shin M, Ferguson M, Willms R, Jones L, Petkau K, Foley E (2022) Immune regulation of intestinal-stem-cell function in Drosophila. Stem Cell Reports 17: 741–755

Singhal R, Shah YM (2020) Oxygen battle in the gut: Hypoxia and hypoxia-inducible factors in metabolic and inflammatory responses in the intestine. J Biol Chem 295: 10493–10505

Solanki S, Shah YM (2024) Hypoxia-Induced Signaling in Gut and Liver Pathobiology. Annu Rev Pathol 19: 291–317

Song Z, McCall K, Steller H (1997) DCP-1, a Drosophila cell death protease essential for development. Science 275: 536–540

Staley B, Irvine K (2010) Warts and Yorkie Mediate Intestinal Regeneration by Influencing Stem Cell Proliferation. Current Biology 20: 1580–1587

Swick L, Kazgan N, Onyenwoke R, Brenman J (2013) Isolation of AMP-activated protein kinase (AMPK) alleles required for neuronal maintenance in Drosophila melanogaster. Biology Open 2: 1321–1323

Takashima S, Hartenstein V (2012) Genetic control of intestinal stem cell specification and development: a comparative view. Stem cell reviews 8: 597–608

Takeishi A, Kuranaga E, Tonoki A, Misaki K, Yonemura S, Kanuka H, Miura M (2013) Homeostatic epithelial renewal in the gut is required for dampening a fatal systemic wound response in Drosophila. Cell Rep 3: 919–930

Tamamouna V, Rahman MM, Petersson M, Charalambous I, Kux K, Mainor H, Bolender V, Isbilir B, Edgar BA, Pitsouli C (2021) Remodelling of oxygen-transporting tracheoles drives intestinal regeneration and tumorigenesis in Drosophila. Nat Cell Biol 23: 497–510

Taylor CT, Colgan SP (2017) Regulation of immunity and inflammation by hypoxia in immunological niches. Nat Rev Immunol 17: 774–785

Trubin S, Patel DB, Tian A (2024) Regulation of the Intestinal Stem Cell Pool and Proliferation in Drosophila. Cells 13

Villareal L, Xue X (2024) The emerging role of hypoxia and environmental factors in inflammatory bowel disease. Toxicological Sciences 198: 169–184

Wang L, Cai Y, Garssen J, Henricks PAJ, Folkerts G, Braber S (2023) The Bidirectional Gut-Lung Axis in Chronic Obstructive Pulmonary Disease. Am J Respir Crit Care Med 207: 1145–1160

Wang L, Zeng X, Ryoo H, Jasper H (2014) Integration of UPRER and Oxidative Stress Signaling in the Control of Intestinal Stem Cell Proliferation. Plos Genetics 10

Wang P, Hou SX (2010) Regulation of intestinal stem cells in mammals and Drosophila. J Cell Physiol 222: 33–37

White K, Grether ME, Abrams JM, Young L, Farrell K, Steller H (1994) Genetic control of programmed cell death in Drosophila. Science 264: 677–683

White K, Tahaoglu E, Steller H (1996) Cell killing by the Drosophila gene reaper. Science 271: 805–807

Wu J, Luo L (2006) A protocol for mosaic analysis with a repressible cell marker (MARCM) in Drosophila. Nature Protocols 1: 2583–2589

Wu M, Yiang G, Liao W, Tsai A, Cheng Y, Cheng P, Li C, Li C (2018) Current Mechanistic Concepts in Ischemia and Reperfusion Injury. Cellular Physiology and Biochemistry 46: 1650–1667

Xie H, Simon MC (2017) Oxygen availability and metabolic reprogramming in cancer. J Biol Chem 292: 16825–16832

Xu N, Wang SQ, Tan D, Gao Y, Lin G, Xi R (2011) EGFR, Wingless and JAK/STAT signaling cooperatively maintain Drosophila intestinal stem cells. Dev Biol 354: 31–43

Zhang P, Edgar B (2022) Insect Gut Regeneration. Cold Spring Harbor Perspectives in Biology 14

Zheng L, Kelly CJ, Colgan SP (2015) Physiologic hypoxia and oxygen homeostasis in the healthy intestine. A Review in the Theme: Cellular Responses to Hypoxia. Am J Physiol Cell Physiol 309: C350–360

Zhou D, Haddad GG (2013) Genetic analysis of hypoxia tolerance and susceptibility in Drosophila and humans. Annual review of genomics and human genetics 14: 25–43

Zhou F, Rasmussen A, Lee S, Agaisse H (2013) The UPD3 cytokine couples environmental challenge and intestinal stem cell division through modulation of JAK/STAT signaling in the stem cell microenvironment. Dev Biol 373: 383–393

Zhou J, Boutros M (2023) Intestinal stem cells and their niches in homeostasis and disease. Cells & Development 175

Zwick RK, Ohlstein B, Klein OD (2019) Intestinal renewal across the animal kingdom: comparing stem cell activity in mouse and Drosophila. Am J Physiol Gastrointest Liver Physiol 316: G313–G322

